# Heterogeneity of Neuronal Populations Within Columns of Primate V1 Revealed by High-Density Recordings

**DOI:** 10.1101/2020.12.22.424048

**Authors:** Shude Zhu, Ruobing Xia, Xiaomo Chen, Tirin Moore

**Author notes:** **Corresponding Author**: Tirin Moore, Dept. of Neurobiology, Fairchild Bldg. 299 Campus Drive, Stanford University School of Medicine, Stanford CA 94305.

## Abstract

Primary visual cortex (V1) has been the focus of extensive neurophysiological investigation, and its laminar organization provides a key exemplar of the functional logic of neocortical microcircuits. Using newly developed high-density, linear array probes, we measured visual responses from large populations of simultaneously recorded neurons distributed across layers of macaque V1. In single recordings, myriad differences in the functional properties of neuronal subpopulations could be observed. In particular, we found that although standard measurements of orientation selectivity yielded only minor differences between laminar compartments, decoding of stimulus orientation from layer 4C responses was superior to that of both superficial and deep layers within the same cortical column. The superior orientation discrimination within layer 4C was associated with greater response reliability of individual neurons rather than lower correlated activity with neuronal populations. The results demonstrate the utility of high-density electrophysiology in revealing the functional organization and network properties of neocortical microcircuits in single experiments.

## Introduction

Early neurophysiological investigations of primary visual cortex (V1) identified the striking emergence of shape processing by orientation-selective neocortical neurons as observed first in the cat (Hubel & Wiesel, 1962) and subsequently in primates (Hubel & Wiesel, 1968). Input from the dorsal lateral geniculate nucleus (dLGN) is fundamentally transformed in V1 from circular, center-surround receptive fields (RFs) into selectivity for orientation in simple cells. A vast number of past studies have examined the distribution of orientation and other functional properties of V1 neurons across cortical layers and morphological cell types in an effort to fully understand the transformation of visual information carried out at this crucial stage of visual processing (Bauer, Dow, & Vautin, 1980; Gur, Kagan, & Snodderly, 2005; Hawken & Parker, 1984; Livingstone & Hubel, 1984; Poggio, Doty, & Talbot, 1977; Ringach, Hawken, & Shapley, 1997; Ringach, Shapley, & Hawken, 2002; Schiller, Finlay, & Volman, 1976). To date, although much is understood about the functional organization and microcircuitry of primate V1, a number of key questions remain unresolved. For example, contrary to early evidence of a lack of orientation selectivity in V1 input layers (4Cα and 4Cβ) (Hubel & Wiesel, 1968), a number of other studies demonstrated that orientation selectivity is more broadly distributed across layers (Hawken & Parker, 1984; Ringach et al., 2002; Schiller et al., 1976). Although a wealth of computational models has been proposed to explain the emergence of orientation selectivity in V1 (e.g. Adorjan, Levitt, Lund, & Obermayer, 1999; Chariker, Shapley, & Young, 2016; McLaughlin, Shapley, Shelley, & Wielaard, 2000), the validity of such models rests on the availability of sufficient data to test key predictions and assumptions.

Historically, the bulk of neurophysiological measurements of the visual properties of macaque V1 neurons have been carried out in successive extracellular recordings from individual neurons or small numbers of neurons using conventional single-electrodes (e.g. Hawken & Parker, 1984; Livingstone & Hubel, 1984; Ringach et al., 1997; Schiller et al., 1976) or low-channel count linear arrays (Hansen, Chelaru, & Dragoi, 2012; Nigam, Pojoga, & Dragoi, 2019; Ziemba et al., 2019). Typically, from such data, the distributions of those properties are studied in aggregated sets of recordings accumulated across multiple sessions. As a result, direct comparisons between subpopulations of neurons within local circuits, e.g. within single cortical columns, are less than ideal. Recent advances in recording technology have facilitated the development of high-density micro-electrode arrays resulting in a substantial increment (~20x) in the number of neurons that can be studied simultaneously within a localized area of neural tissue. A prime example is the recent development of the Neuropixels probe (IMEC, Inc.), which consists of a high-channel count Si shank with continuous, dense, programmable recording sites (~1000/cm). Numerous recent studies have demonstrated the advantages of such probes, such as their use in recording large neuronal populations within deep structures where optical approaches cannot be deployed (Jun et al., 2017; Steinmetz, Zatka-Haas, Carandini, & Harris, 2019). However, only a few electrophysiological studies of the primate brain have been carried out thus far (Hesse & Tsao, 2020; Trautmann et al., 2019), and none have targeted primate V1.

Using Neuropixels probes, we studied the visual activity of populations of neurons distributed across layers of macaque V1. The large capacity of Neuropixels probes facilitated comparisons between substantial populations of neurons within single cortical columns, both within and between defined laminar compartments. Robust differences in the functional properties of neurons within different subpopulations were observable within a single recording session. For example, synchrony and spike count correlations among layer 4C neurons differed dramatically from those of superficial and deep layer neurons. Most surprisingly, although standard measurements of orientation selectivity yielded only minor differences between laminar compartments, we found that decoding of orientation from layer 4C neuronal responses was superior to that of superficial and deep layer neurons within the same cortical column. Furthermore, the superior orientation decoding from layer 4C activity was associated with greater response reliability of individual neurons rather than from the lower correlated activity among layer 4C neurons. The results demonstrate the utility of high-density electrophysiology in revealing the functional organization and network properties of primate neocortical microcircuits in single experiments.

## Results

We recorded the activity of neurons in V1 of 2 anesthetized monkeys (M1, M2) using high-density, multi-contact Neuropixel probes (version 3A; IMEC Inc, Belgium)(Figure 1A) (METHODS). Each probe consisted of 986 contacts (12 μm x 12 μm, 20 μm spacing) distributed across 10 mm, of which 384 contacts could be simultaneously selected for recording. Probes were inserted into the lateral operculum of V1 with the aid of a surgical microscope at angles nearly perpendicular to the cortical surface. The dense spacing between electrode contacts provided multiple measurements of the waveforms from individual neurons (mean = 4.52) and facilitated the isolation of each of a large number of single neurons (Figure 1B), typically >300 in a single penetration. In total, we recorded the activity of 1,833 well-isolated single neurons across layers of V1 in 5 penetrations in the two monkeys (1,124 neurons, M1; 709 neurons, M2). In each of the 5 recordings, we studied the functional properties of simultaneously recorded populations of neurons within different laminar compartments. In order to assess the distribution of properties of V1 neurons across layers, we first estimated the borders of laminar compartments by combining the histological data with current-source density (CSD) measurements (Fig. 1C) in each recording (METHODS). Using those estimates, we assigned each of the recorded neurons to a specific laminar compartment. Cortical layers were divided into four comparably sized laminar compartments, specifically layers 2/3, 4A/B, 4C, and 5/6 (Mean depth: 650μm, 311μm, 281μm, 489μm, respectively). Notably, we combined layers 4Cα and 4Cβ, respectively the Magnocellular and Parvocellular recipient layers, into a single compartment in order to achieve comparable numbers of recorded neurons in each compartment.

**Figure 1.**
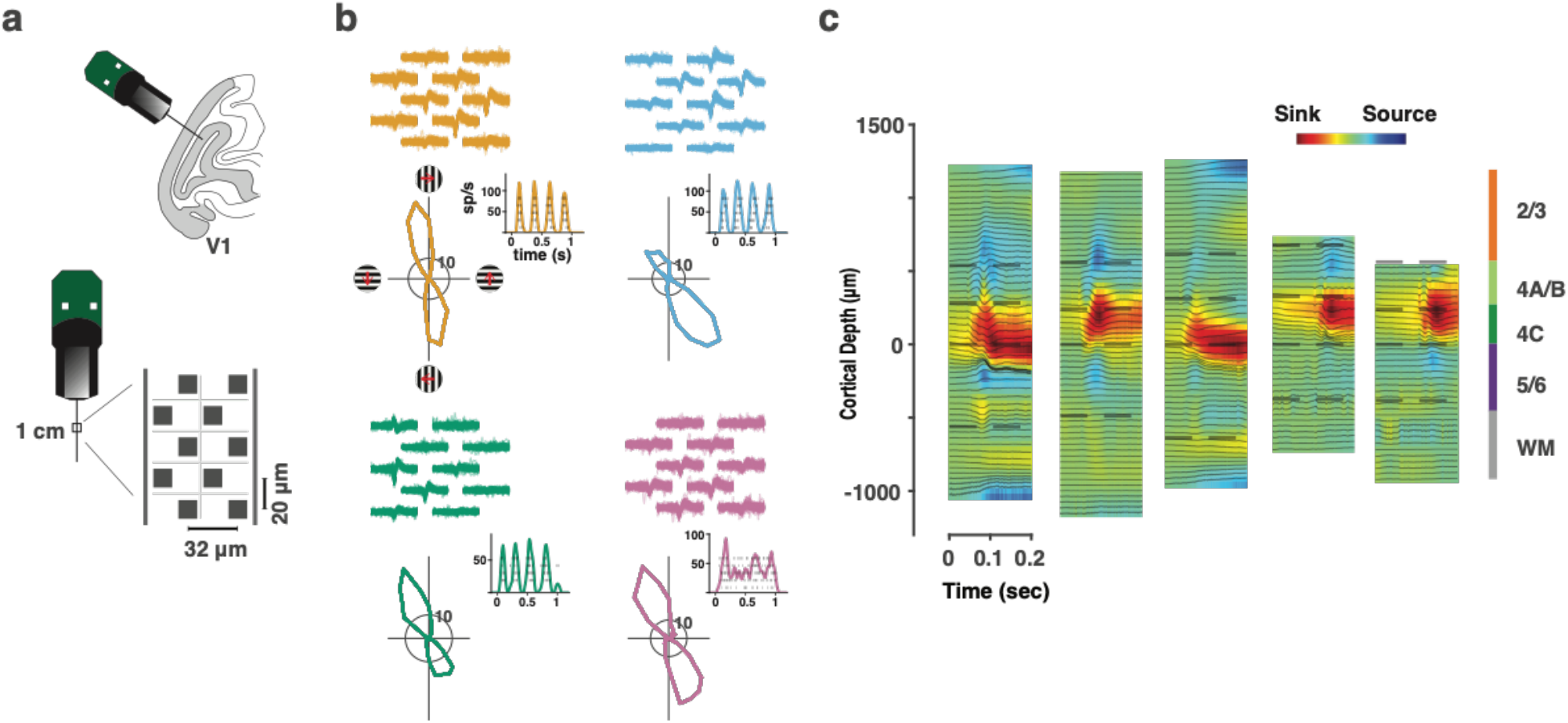
Neuropixels recordings in primate V1. **a,** Upper cartoon depicts the angle of probe penetrations made into the lateral surface and underlying calcarine sulcus of V1. Lower, Image of Neuropixels probe base and shank. Right diagram shows the layout of electrode contacts for a section of the recording shank. **b**, Example single-neuron recordings with Neuropixels probes, three simple cells (orange, blue, green) and 1 complex cell (purple). Top, neuronal waveforms recorded across multiple adjacent electrode contacts are shown for each neuron. Bottom, each neuron’s response to its preferred orientation (rasters and instantaneous spike rates) and their corresponding tuning curves. Red arrows denote the drift direction of oriented gratings. **c,** CSD profiles for each of the five recording sessions. CSDs were derived from LFP responses to drifting gratings. In each session, laminar compartment boundaries (dashed lines) were determined using histological data and the CSD profile.

### Differential distribution of neuronal waveforms across V1 compartments

First, we found that neurons with different waveform shapes were distributed unequally across cortical layers. As expected, extracellular spike waveforms throughout most of the recording depths typically exhibited initial negative components, followed by positive ones of variable length (Fig. 2A). At the deepest recording sites, the positive and negative components of spike waveforms were reversed, characteristic of axonal spikes (Schomburg, Anastassiou, Buzsaki, & Koch, 2012). Accordingly, we labeled these latter waveforms as putative ‘axonal’ spikes (Fig. 2B). We compared the frequencies of putative axonal waveforms between white matter (Wm) and all combined gray matter (Gm) compartments using the laminar boundaries derived from the CSD and histological data. These comparisons revealed that the density of axonal spikes (n per 100 ***μ***m) was significantly larger within the Wm than in the Gm (Median Wm: 6.9, Gm 2.2, p=0.0159, Mann-Whitney U test) (Fig. 2C). Moreover, axonal waveforms within the Wm were much more frequent than other waveform types in both animals (M1, 7% Gm, 60% Wm, p=5.7E^−56^; M2, 17% Gm, 72% Wm, p=5.3E^−39^, ***χ***^2^ test). This result thus corroborated our estimates of Gm-Wm boundaries.

**Figure 2.**
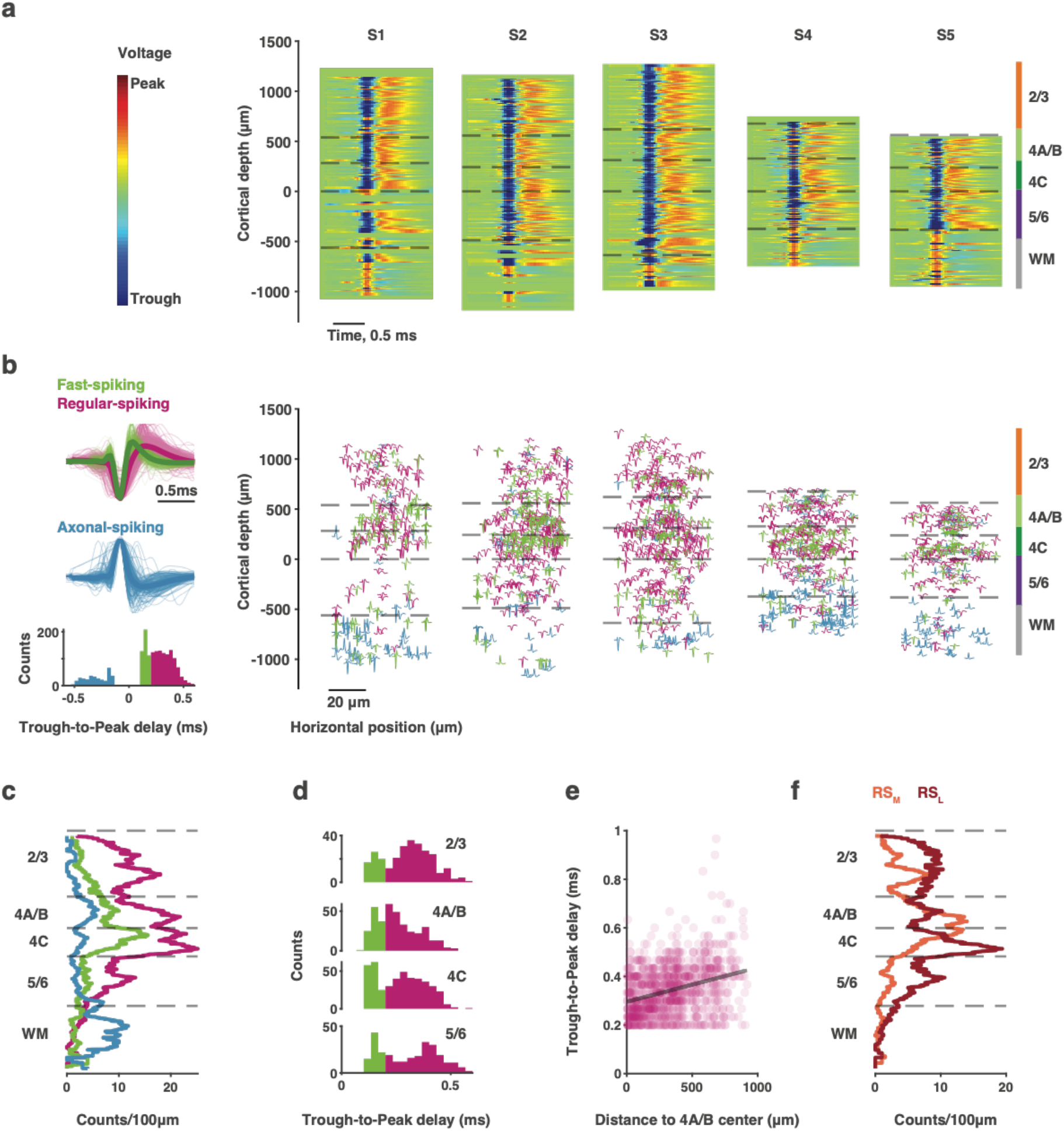
Waveform classes and their laminar distributions. **a,** Spike waveform heat maps of neurons recorded across cortical depth in all five sessions. Laminar compartments are indicated as in Fig. 1. **b,** Top left, three waveform classes identified from their trough-to-peak delay, regular-spiking, fast-spiking, and putative axonal (negative delay). Bottom left, histogram of trough-to-peak delays for all neurons. Right, the location of neurons within different waveform classes across cortical depth (and horizontal position). Horizontal axes are magnified for visualization. **c,** Density of waveform classes across cortical depth for all sessions combined. Densities were computed within a sliding 100-μm window. **d,** Histogram of trough-to-peak delays for neurons within each laminar compartment for all sessions combined. **e,** Trough-to-peak delays of RS neurons increase with distance from Layer 4A/B. **f**, Density of two subclasses of RS waveforms across cortical depth.

A number of past studies have exploited known differences in somatic spike waveform durations to potentially distinguish different functional classes of cortical neurons in extracellular recordings (McCormick, Connors, Lighthall, & Prince, 1985; Mitchell, Sundberg, & Reynolds, 2007; Mountcastle, Talbot, Sakata, & Hyvarinen, 1969; Wilson, O’Scalaidhe, & Goldman-Rakic, 1994), specifically so-called regular-spiking (RS) and fast-spiking (FS) neurons. These two classes correspond respectively to putative excitatory neurons and inhibitory interneurons, though with important exceptions (Kawaguchi & Kubota, 1997). Thus, we classified all non-axonal spike waveforms initially as RS or FS, according to standard criteria (Methods)(Fig. 2B-C). As expected, RS neurons outnumbered FS neurons roughly 2.4:1 (RS: 963; FS: 406). For both types of neurons, the density of neurons varied significantly across layers (RS: p = 0.0167; FS: p = 0.0093. Kruskal-Wallis test), with greater numbers of both types in Layer 4C (Supplementary Table 1). However, the disproportionality of the two waveform classes differed significantly across cortical layers (***χ***=9.2010, df=3, p=0.02673, ***χ***^2^ test), due to more equal proportions (~2:1) of cell types within layers 4A/B and 4C.

In addition to the two major subclasses of somatic waveforms, we found that the trough-to-peak durations of RS neurons varied widely across laminar compartments (Fig. 2D). In particular, durations were greater within supragranular and infragranular layers. Overall, waveform durations of RS neurons increased with distance from Layer 4A/B (r=0.32, p=1.74E^−26^)(Fig. 2E). This suggests that perhaps at least one additional subclass of broader spiking neurons exists within V1 and is generally consistent with evidence of more than two distinct classes of spike waveforms within mammalian neocortex (e.g. Munoz, Tremblay, & Rudy, 2014; Trainito, von Nicolai, Miller, & Siegel, 2019). Thus, we separated the two putative subclasses of RS neurons accordingly into Regular-medium (RS_M_) and long-broad (RS_L_) (Fig. 2D). The densities of the two classes peaked at different cortical depths; RS_L_ neurons peaked at the 4C-5/6 border, and RS_M_ neurons peaked within the 4A/B compartment. Consequently, the distributions of the two subclasses of RS neurons differed across layers (***χ***=64.3571, df=3, p=6.88E^−14^, ***χ***^2^ test)(Fig. 2F).

### Differences in correlated activity between laminar compartments and waveform types

The high-density recordings also allowed us to measure correlated activity among many 1000s of neuronal pairs within and between layers and neuronal subtypes, thus providing a robust assay of network connectivity within single recording sessions. We first measured synchrony (cross-correlograms) and spike count (‘noise’) correlations in the responses of neuronal pairs to visual stimulation (Methods). The two measures gauge the degree of correlated spiking activity, but at short and long timescales, respectively (Averbeck, Latham, & Pouget, 2006; Cohen & Kohn, 2011). Both measures were computed from the responses of 3,486-17,044 neuronal pairs in each recording session. Cross-correlograms (CCGs) measure the pairwise correlation structure of spiking activity, the peak amplitude of which is thought to gauge the strength of the functional connectivity between two neurons (Alonso & Martinez, 1998; Perkel, Gerstein, & Moore, 1967b; Toyama, Kimura, & Tanaka, 1981) (Fig. 3A). The temporal delay of the CCG peak indicates the lead-lag relationship between the activity of neuronal pairs. We computed CCGs for all pairs of neurons using their spike trains evoked during visual stimulation. Pairs of neurons were drawn from a total of 676 cells (461 M1, 215 M2) with significant visual responses to drifting sinusoidal gratings. In most of the recording sessions, data were obtained from >17 visually responsive neurons recorded in each compartment (Supplementary Table 2). Figure 3B shows the network of CCGs for all neuronal pairs across cortical depth for a single session (S3). The CCG network not only reveals apparent local connections between neurons within the same laminar compartment (e.g. layer 2/3), but also a large number of strong distant connections between neurons across layers. In addition, the network revealed a greater tendency of layer 2/3 neurons to lag responses of neurons in all other layers.

**Figure 3.**
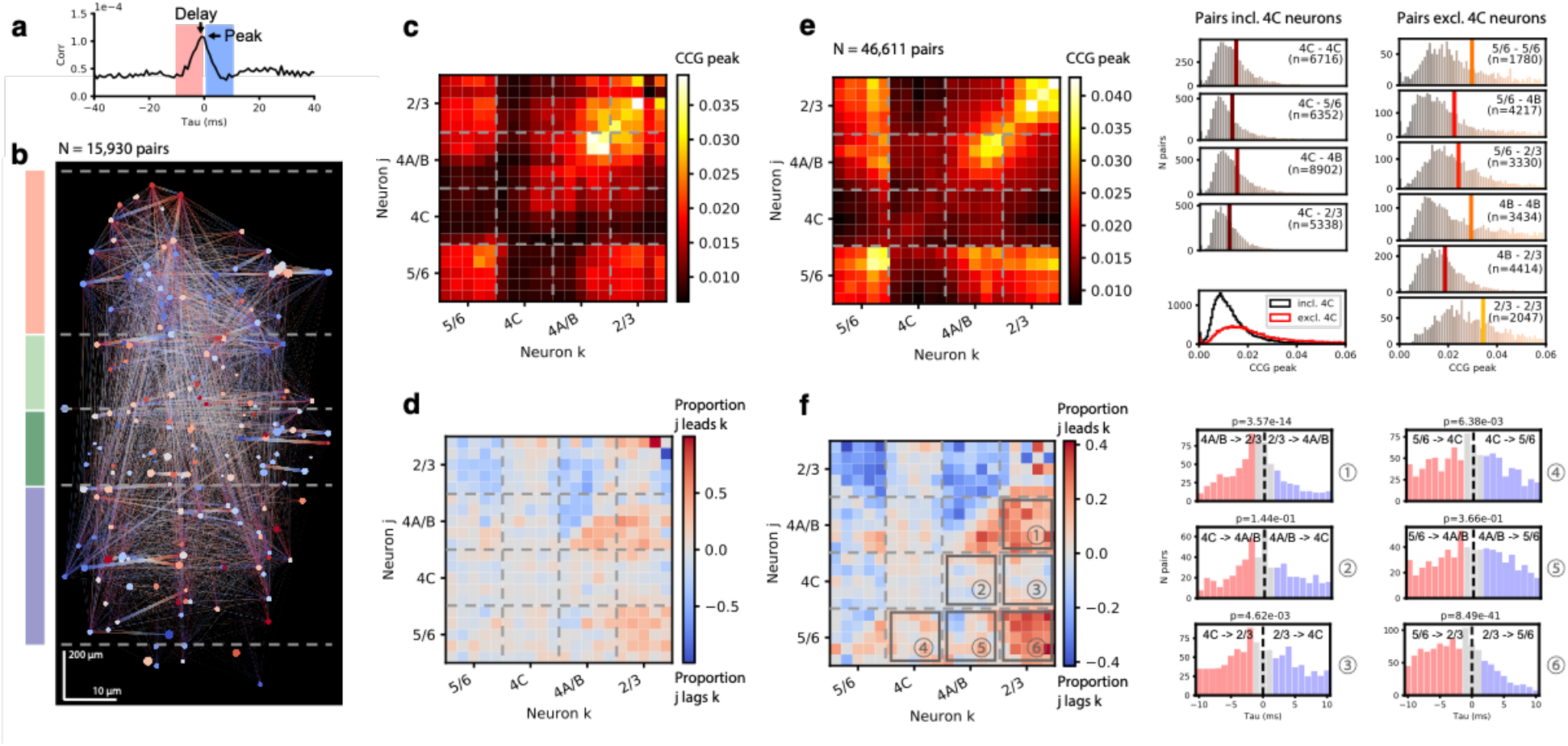
Cross-correlations in neuronal activity across V1 layers. **a**, An example CCG of a single pair of neurons. Arrows denote the CCG peak and the CCG delay (tau at the peak). **b,** CCG network diagram from session S3. Dots mark individual neurons at their estimated cortical positions. Connecting lines denote the CCG peak (thickness) and the proportion of lead-lag relationship (direction of color gradient). Dot size and color respectively denote the mean of CCG peaks and proportion of lead-lag relationships for all pairings of each neuron. **c**, Matrix of CCG peaks across cortical depth from session S3. Each pixel shows the mean CCG peak across neuronal pairs at each depth. Cortical depth is aligned to the 4C - 5/6 border and normalized by the thickness of each compartment. **d**, Correlation direction matrix across cortical depth from session S3. Each pixel shows the proportion of lead-lag relationships between neuronal pairs. **e**, CCG peak matrix for all sessions combined. Right columns show the distributions of peaks for neuronal pairs within and between laminar compartments that include (left) or exclude (right) layer 4C neurons, and a comparison of the two (bottom left). **f**, Correlation direction matrix of all sessions combined. Right columns show the distribution of CCG delays for neuronal pairs between laminar compartments.

For the same session, the average peak correlations for neuronal pairs were compiled into a matrix of correlations across cortical depth (Fig. 3C). This matrix revealed measurable synchrony among neurons within and between all layers, with the exception of 4C. Synchrony was notably weak between 4C pairs, as were distant pairings involving 4C neurons. Overall, correlations between pairs excluding layer 4C neurons, either within or between laminar compartments, were significantly greater than those including 4C neurons (t-test, p=5.87E^−178^), and those differences highlighted the boundaries between assigned laminar compartments. These differences were not a result of differences in mean firing rate between the different laminar compartments (Fig. S1A). In addition, similar to the pattern observed in the CCG network (Fig. 3B), layer 2/3 neurons were significantly more likely to lag neurons within other layer compartments (sign test, 5/6 to 2/3: p=8.49E^−41^; 4C to 2/3: p=4.62E^−3^; 4A/B to 2/3: p=3.57E^−14^) (Fig. 3D). Both results were similar in the remaining 4 sessions (Fig. S2). Analyses of data combined across sessions revealed significantly lower synchrony for layer 4C neurons (t-test, p<1.00E^−200^)(Fig. 3E), and showed that layer 2/3 neurons tended to lag neurons in other layers (sign test, 5/6 to 2/3: p=8.06E^−60^; 4C to 2/3: p=1.97E^−4^; 4A/B to 2/3: p=4.84E^−73^)(Fig. 3F). Both of these major observations are consistent with previous evidence. Noise correlations were previously shown to be significantly lower in the input layers (Hansen et al., 2012). The lag in layer 2/3 is consistent with an overall convergence of inputs toward layer 2/3 neurons from which projections to extrastriate areas largely originate (Callaway, 1998), though circuit models of V1 often posit deep layers as a later stage corticofugal output (Gilbert & Wiesel, 1983).

The pattern of differences in overall synchrony observed across laminar compartments was similar for comparisons of noise correlations; neurons in layer 4C exhibited considerably lower noise correlations than more superficial or deep compartments (Fig. 4A, Fig. S3) (t test, p<1.00E^−200^). In addition, we measured correlation coefficients between responses to different stimuli (i.e., signal correlations) in order to quantify the extent to which pairs of neurons exhibited similar tuning properties. As with synchrony and noise correlations, overall mean signal correlations differed significantly across laminar compartments (ANOVA, p=7.42E^−30^). However, the pattern of results was reversed. Rather than exhibiting lower correlations, signal correlations in layer 4C were greater than those of layers 5/6 and 4A/B (5/6: mean=0.307, 4C: mean=0.420, 4A/B: mean=0.314; Post Hoc t test, 5/6 vs. 4C: p=1.94E^−15^, 4A/B vs. 4C: p=1.28E^−20^) and were comparable with layer 2/3 (2/3: mean=0.423, 2/3 vs. 4C: p=0.80). Furthermore, layer 5/6 neurons, from which the highest levels of synchrony and noise correlations were measured, exhibited lower signal correlations than compartments 4C and 2/3 (Post Hoc t test, 5/6 vs. 4C: p=1.94E^−15^, 5/6 vs. 2/3: p=5.19E^−17^). As with synchrony, these differences did not result from differences in mean firing rate between the different laminar compartments (Fig. S1B).

**Figure 4.**
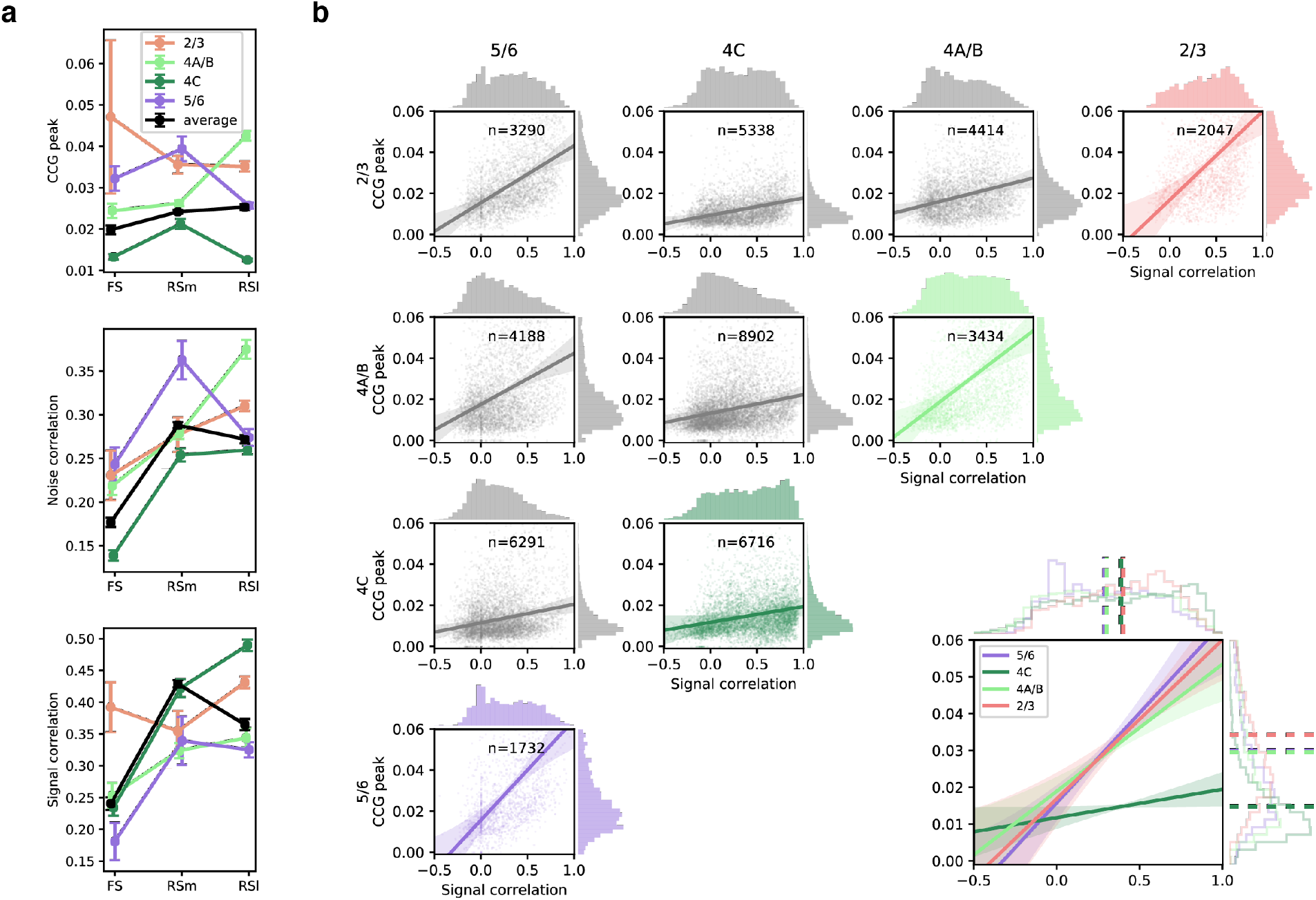
Synchrony, noise, and signal correlations across laminar compartments and waveform classes. **a,** Mean CCG peaks, noise, and signal correlations within different compartments and for different neuronal waveform classes. Error bars denote +/− S.E.M.. **b,** Relationship between CCG peak and signal correlation for all neuronal pairs within and between laminar compartments. Right plot shows mean CCG peak - signal correlation relationship for pairs within each compartment. Lines denote linear fits; shaded region shows +/− S.E.M.. Dotted vertical and horizontal lines denote marginal means.

Next, we compared synchrony, noise and signal correlations across the 3 waveform classes (FS, RS_M_, and RS_L_). Overall, there was a main effect of waveform class on all 3 measures (ANOVA, synchrony: p=0.03; noise correlation: 6.16E^−54^; signal correlation: p=1.27E^−54^). Moreover, significant differences were observed for all pairwise comparisons between classes for each measure (Supplementary Tables S3-5). Within a given laminar compartment, measures of synchrony and correlation could differ between waveform classes by more than a factor of 2. Notably, lower synchrony, noise, and signal correlations were generally exhibited by FS neurons. These differences were not a result of differences in mean firing rate between the different neuron classes (Fig. S4). Thus, synchrony, noise, and signal correlations among neurons clearly depended on both layer and waveform type.

We next assessed the dependence of synchrony on the similarity of tuning properties between neuronal pairs across the different laminar compartments. Previous studies have shown that greater functional and synaptic connectivity typically occurs between neurons with similar stimulus preferences (Cossell et al., 2015; Denman & Contreras, 2014; Lee et al., 2016). Thus, we hypothesized that the strength of synchrony between V1 neurons should depend on the similarity of visual properties of neurons within the same cortical column, i.e. their signal correlations. Moreover, we wondered if the lower synchrony values observed among layer 4C neurons might be accompanied by correspondingly low signal correlations. For virtually all neuronal pairings, both within and between laminar compartments, we observed a positive relationship between CCG peaks and signal correlations (p<10^−30^ for all linear regressions) (Fig. 4B). However, the relationships differed across laminar compartments. Specifically, within layer 4C, synchrony was dramatically less dependent on signal correlations (slope t test, 4C vs. 5/6: p=4.79E^−45^; 4C vs. 4A/B: p=4.53E^−63^; 4C vs. 2/3: p=3.72E^−26^), such that even at high levels of signal correlation (i.e. >0.5), synchrony remained very low. Thus, layer 4C neurons exhibited a qualitatively different relationship between the signal correlation and synchrony. The limited functional connectivity among layer 4C neurons was present in spite of substantial signal correlations, and thus strong similarities in stimulus preference.

### Visual properties of simultaneously recorded neurons across V1 layers

A wealth of past electrophysiological studies has explored the differences in the functional properties of neurons across the layers of primate V1 (Bauer et al., 1980; Gur et al., 2005; Hawken & Parker, 1984; Livingstone & Hubel, 1984; Poggio et al., 1977; Ringach et al., 2002; Schiller et al., 1976). The most classically examined properties include firing rates, the proportions of simple and complex cells, the incidence of direction selectivity, and various components of orientation selectivity. The high density recordings enabled us to assess these properties in the large numbers of visually responsive neurons recorded simultaneously in single sessions. Drifting circular Gabor gratings were presented for 1 second within the joint RFs of recorded neurons at 36 different directions (0 – 360°, 10° step). Four spatial frequencies (0.5, 1, 2, 4 cycle/deg.) were tested and responses to the optimal spatial frequency were used in the analyses (Methods). Consistent with previous studies, we found significant difference in the maximum firing rates of neurons located across laminar compartments (Kruskal-Wallis test, ***χ***^2^(3) =16.65, *P*=.0008), with the highest median rates found in layer 4C (Fig. 5A). Next, we compared the distribution of simple and complex cells across layers. Simple and complex cells are known to differ dramatically in their response to drifting gratings in that simple cells, being sensitive to phase, exhibit robust oscillatory modulation, while complex cells do not (De Valois, Albrecht, & Thorell, 1982) (Figure 1B). Thus, we could use the modulation ratio of visual responses to reveal any differential distribution of simple and complex cells across layers in each recording (Figures 5B). As expected, modulation ratios varied significantly across laminar compartments (Kruskal-Wallis test, ***χ***^2^(3) =28.55, *P*=2.79E^−6^), with larger ratios found among layer 4C neurons.

**Fig 5.**
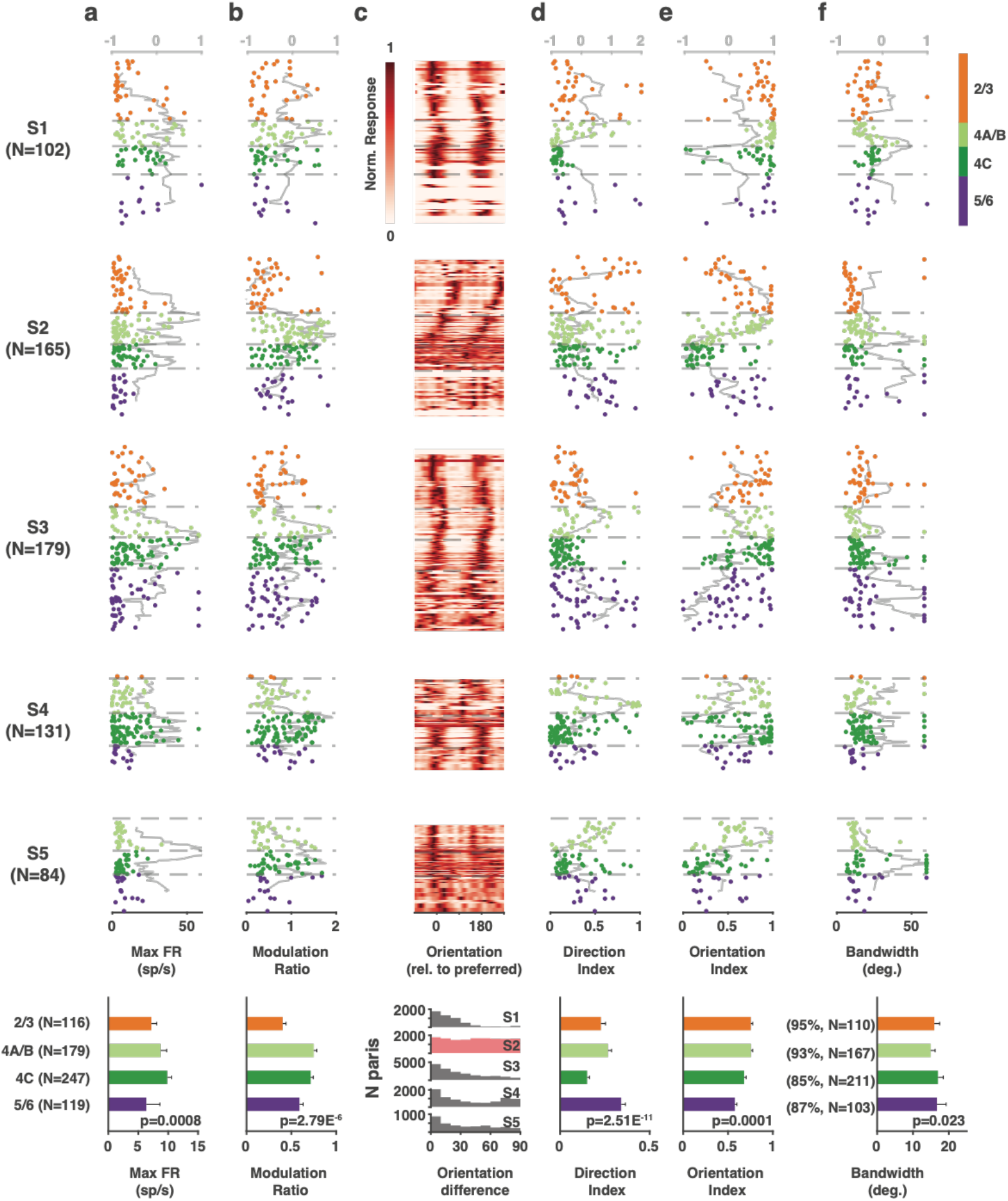
Functional properties of single V1 neurons recorded simultaneously across laminar compartments. Components of visual responses of all V1 neurons recorded in five sessions across identified laminar compartments. **a**, Maximum firing rate response (to preferred stimulus). **b**, Modulation ratio. **c**, Heat map of visual responses across drift direction of oriented grating. (vertical thickness is greater for less dense neuronal populations.) **d**, Direction index. **e**. Orientation index. **f**. Orientation tuning bandwidth. Data from each neuron is plotted at its corresponding cortical depth. Gray lines (top abscissa) denote kurtosis of each value in a running 100 μm window. Bottom row plots show averaged results across all 5 sessions.

Given that our probe penetrations were made largely perpendicular to the cortical surface, we could visualize the known columnar organization of orientation tuning in V1 by simply plotting the orientation preference of individual neurons recorded across the cortical depth. Gratings drifted across all directions, and a majority of neurons exhibited peak responses at two orientations that were equal but moving in opposite directions (Fig. 5C). Moreover, for most recordings (sessions 1,3-5), the preferred orientation remained similar for neurons distributed across depth, indicating that these penetrations remained largely within a single orientation column (*columnar* sessions). In contrast, the preferred orientation varied systematically across cortical depth in session 2 (*non-columnar* session). Across recordings, a subset of neurons responded more strongly to one of the drift directions, thus exhibiting direction selectivity. Direction selectivity was quantified using a standard selectivity index that compared the preferred drift direction to the opposite direction (Methods) (Figures 5D). Direction selectivity varied significantly across layers (Kruskal-Wallis test, ***χ***^2^(3) =52.36, *P*=2.51E^−11^), with significantly lower values in layer 4C (Two-tailed Mann-Whitney U test, *P*=1.55E^−11^). Similarly, for each neuron, we also computed a selectivity index for orientation, by comparing responses to the preferred and orthogonal orientations (Methods)(Figures 5E). As with direction selectivity, orientation selectivity varied significantly across layers (Kruskal-Wallis test, ***χ***^2^(3) =20.33, *P*=.0001). However, consistent with previous studies (Hawken & Parker, 1984; Ringach et al., 2002; Schiller et al., 1976), orientation selectivity was not significantly lower in layer 4C than in other layers (All sessions: *n_4C_*=247, *n_others_*=414, Median: 4C 0.68, others 0.70, *P*=.084; All columnar sessions: Median: 4C 0.77, others 0.72, *P*=.52; Two-tailed Mann-Whitney *U* test). For most of the recorded neurons, responses across orientation were well fit by a circular Gaussian (median *R*^2^ = 0.95)(Methods). From the population of well fit neurons (*R*^2^ ≥ 0.7, 89.4 %), we obtained tuning bandwidths and compared them across cortical depth (Figures 5F). Overall, tuning bandwidths differed significantly across layers (Kruskal-Wallis test, ***χ***^2^(3) =9.51, *P*=.023), consistent with previous evidence (Ringach et al., 2002). Bandwidth was very slightly, but significantly, greater in 4C compared to other layers (All sessions: *n_4C_*=211, *n_others_*=380, Median: 4C 17.0, others 15.7, *P*=.012; All columnar sessions: Median: 4C 17.5, others 16.5, *P*=.011; Two-tailed Mann-Whitney *U* test).

### Superior decoding of orientation from layer 4 neurons

Orientation selectivity emerges within primate V1 and is thus perhaps the most fundamental property of primate V1 neurons. Differences in orientation selectivity of neurons within different layers have been the focus of numerous previous studies (Bauer et al., 1980; Gur et al., 2005; Hawken & Parker, 1984; Livingstone & Hubel, 1984; Poggio et al., 1977; Ringach et al., 1997; Ringach et al., 2002; Schiller et al., 1976). Similar to our observations, these studies found equivocal differences in the orientation selectivity of individual neurons distributed across layers. However, classical measurements of selectivity (e.g. bandwidth, selectivity index) are limited in their ability to adequately quantify the information contained in the responses of sensory neurons. As an alternative, more recent studies have deployed machine learning algorithms to decode orientation signals contained in the responses of populations of simultaneously recorded V1 neurons (Berens et al., 2012; Graf, Kohn, Jazayeri, & Movshon, 2011). These studies capture the rapid and highly orientation-sensitive signals conveyed by populations of V1 neurons. However, recordings in these studies did not allow for simultaneous comparisons of orientation decoding between different subpopulations of neurons within layers of the same cortical column. Thus, we leveraged our high-density recordings to examine the strength of orientation signals within different V1 layers using a decoding approach. Specifically, we employed a Linear-Discriminant Analysis (LDA) decoder (Fisher, 1936; Mendoza-Halliday & Martinez-Trujillo, 2017) to predict visual stimuli based on the combined activity of subpopulations of simultaneously recorded neurons distributed across layers of V1.

LDA decoders were trained to discriminate between pairs of stimulus orientations from the activity of neuronal subpopulations in each recording session (Methods). First, we compared the performance of the decoder at discriminating orientation changes, relative to the peak (preferred) orientation, using responses from a constant number of neurons (n=10) in each laminar subpopulation (Fig 6A). In this comparison, orientation change thresholds (minimal Δθ for performance exceeding 50%) did not differ between laminar compartments, perhaps due to a limited resolution in orientation sampling near the peak orientation (Δθ = 10deg.). However, for the majority of recording sessions, decoders trained on the activity of populations within different laminar compartments exhibited clear differences in orientation discrimination. Specifically, we found that populations of layer 4C neurons consistently outperformed populations in other layers, particularly those within layers 2/3 and 5/6. Only in the non-columnar session, in which the preferred orientation varied widely across cortical depth (Fig. 5C), was the performance of layer 4C decoders inferior to that of other layers. When compared to other laminar subpopulations, the average suprathreshold performance of layer 4C decoders in columnar sessions significantly exceeded that of superficial (L2/3) and deep (L5/6) layers (4C vs 2/3: mean Δ% = 9.9 ± 1.7; 4C vs 5/6: mean Δ%=15.2 ± 2.6; *P*=.0312, Two-tailed Wilcoxon signed-rank test). Decoding performance for layer 4A/B neurons also exceeded that of superficial (L2/3) and deep (L5/6) layers (4A/B vs 2/3: mean Δ% = 9.4 ± 0.1; 4A/B vs 5/6: mean Δ%= 12.0 ± 1.0; *P*=.0312, Two-tailed Wilcoxon signed-rank test). We considered that the apparent superiority of layer 4 neurons at discriminating orientation could have resulted from the arbitrary number of neurons chosen (n=10) in each neuronal subset. Thus, for each session, we also generated neuron-dropping curves (NDCs) (Wessberg et al., 2000) from the performance of neuronal subsets obtained in each laminar compartment in order to compare performances across varying population sizes (Fig. 6B). For each of the columnar sessions, the NDC revealed greater performance for layer 4 neurons (4A/B and 4C) across the range of population sizes compared to supragranular and infragranular compartments.

**Fig 6.**
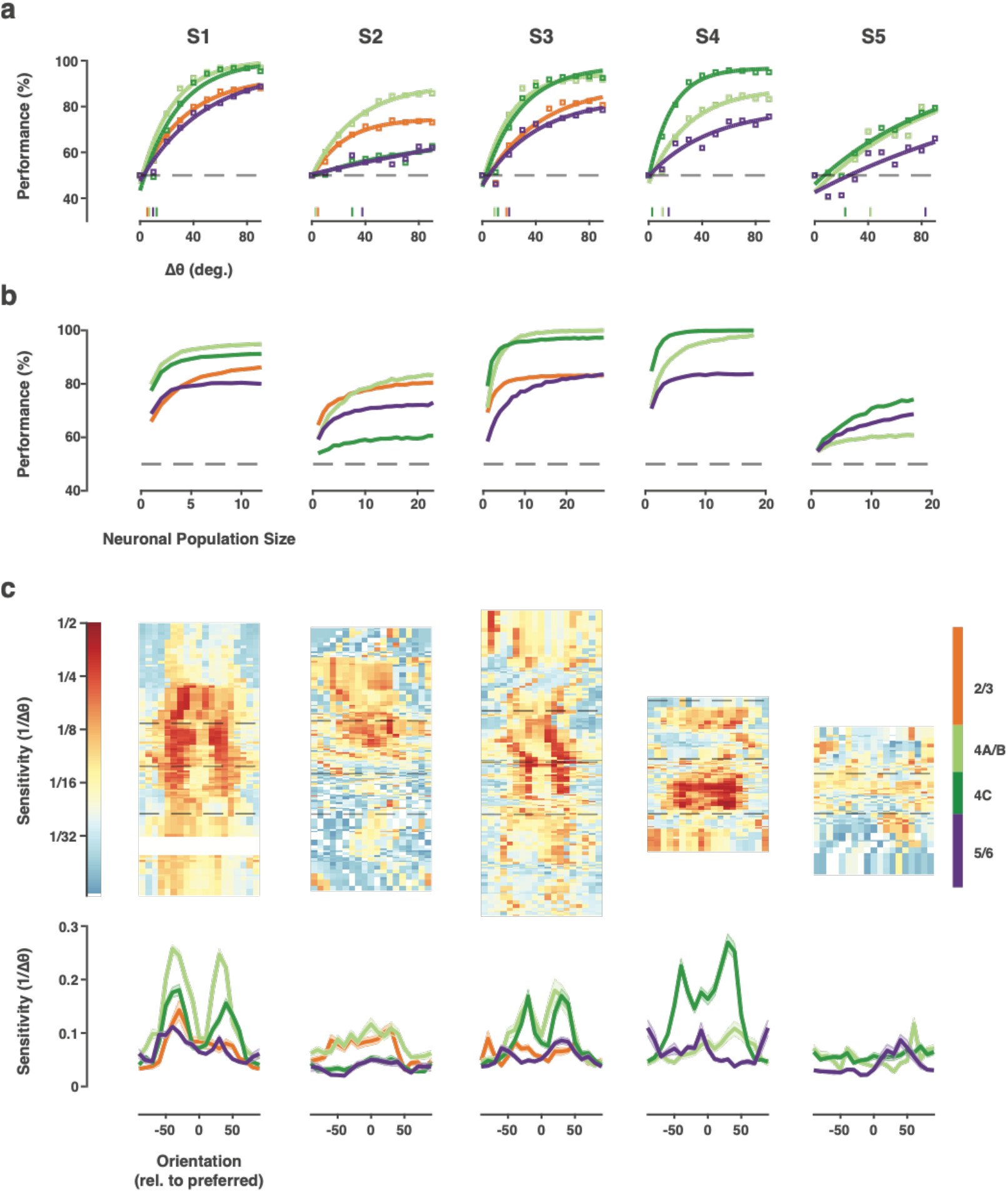
Decoding of orientation from laminar subpopulations of V1 neurons. **a,** Performance of decoders trained to discriminate between the preferred and a second orientation. Decoders were trained from the responses of populations of neurons within different laminar compartments. Each point shows the average decoder performance using the activity of fixed subsets of 10 neurons within each laminar subpopulation for 9 pairwise orientation discriminations. Lines denote curve fits for each compartment. Results from each recording session are shown separately. Vertical lines on abscissa indicate orientation thresholds. **b,** Neuron dropping curves for the different laminar subpopulations. **c,** Sensitivity of orientation decoding across cortical depth. Each point in the heat map shows the orientation sensitivity (1/ threshold) for subpopulations of neurons based on a 60% performance threshold in decoder discrimination of orientation changes from different comparison points. Each heat map is aligned to the preferred orientation of the recorded cortical column. Bottom plots show average sensitivities across orientation comparison points for each laminar compartment. Error bars denote +/− S.E.M..

Next, we measured the sensitivity to laminar subpopulations to orientation changes across the range of orientations. Measurements of pairwise discrimination performance were obtained for all combinations of the 18 orientations (n=153). For each orientation, sensitivity was measured as the reciprocal of the threshold change in orientation required to exceed 60% performance in subsets of 10 neurons distributed across layers. Peaks in orientation change sensitivity were typically observed on the flanks of the preferred orientation of the constituent neurons (Fig. 6C) corresponding to the steepest points in the orientation tuning curves. Across sessions, sensitivity was consistently highest in middle layers 4A/B and 4C, with the lowest values found in the 5/6 compartment. Thus, in the columnar recordings, populations of layer 4 neurons exhibited greater orientation sensitivity than their superficial and deep layer counterparts.

### Single neuron properties contribute to superior orientation decoding in layer 4C

We next considered the extent to which the superior decoding of orientation in layer 4 might be due to the reduced correlated variability there. Much experimental and theoretical work describes how noise correlations can reduce or limit the amount of information available in the responses of neuronal populations (Abbott & Dayan, 1999; Averbeck et al., 2006; Cohen & Kohn, 2011). Indeed, superior orientation discrimination in layer 4 was previously predicted from the observation of reduced correlated variability (Hansen et al., 2012). However, superior Layer 4 performance appeared to be present even in very small populations, or even single neurons (NDCs, Fig. 6B), suggesting that correlated variability was not a key factor. To address this more directly, we repeated the decoding comparisons using shuffled trials, thus removing correlated activity (Methods). As with the unshuffled datasets, decoders trained on the activity of populations within different laminar compartments exhibited clear differences in orientation discrimination in columnar recordings. In the trial shuffled populations, we also found that populations of layer 4 neurons consistently outperformed populations in layers 2/3 and 5/6 (Fig. 7A). When compared to other laminar subpopulations, the average suprathreshold performance of layer 4C neurons in columnar sessions significantly exceeded that of superficial (L2/3) and deep (L5/6) layers (4C vs 2/3: mean Δ% = 9.8 ± 1.8; 4C vs 5/6: mean Δ%=15.4 ± 2.7; *P*=.0312, Two-tailed Wilcoxon signed-rank test). Decoding performance for layer 4A/B neurons also exceeded that of superficial (L2/3) and deep (L5/6) layers (4A/B vs 2/3: mean Δ% = 9.4 ± 0.2; 4A/B vs 5/6: mean Δ%= 12.3 ± 0.8; *P*=.0312, Two-tailed Wilcoxon signed-rank test). We also directly compared the discrimination performance between shuffled and unshuffled datasets for all pairwise orientation differences (Fig. 7B). These comparisons revealed no differences, or only very small differences, for any of the laminar compartments (2/3: mean Δ%=-0.060 ± 0.025; *P*=.0181; 4C: mean Δ%=-0.001 ± 0.014; *P*=.942; 4A/B: mean Δ%=-0.018 ± 0.015; *P*=.233; 5/6: mean Δ%=-0.066 ± 0.020; *P*=.001. Two-tailed t test), confirming that trial shuffled populations produced the same results as the unshuffled populations.

**Fig 7.**
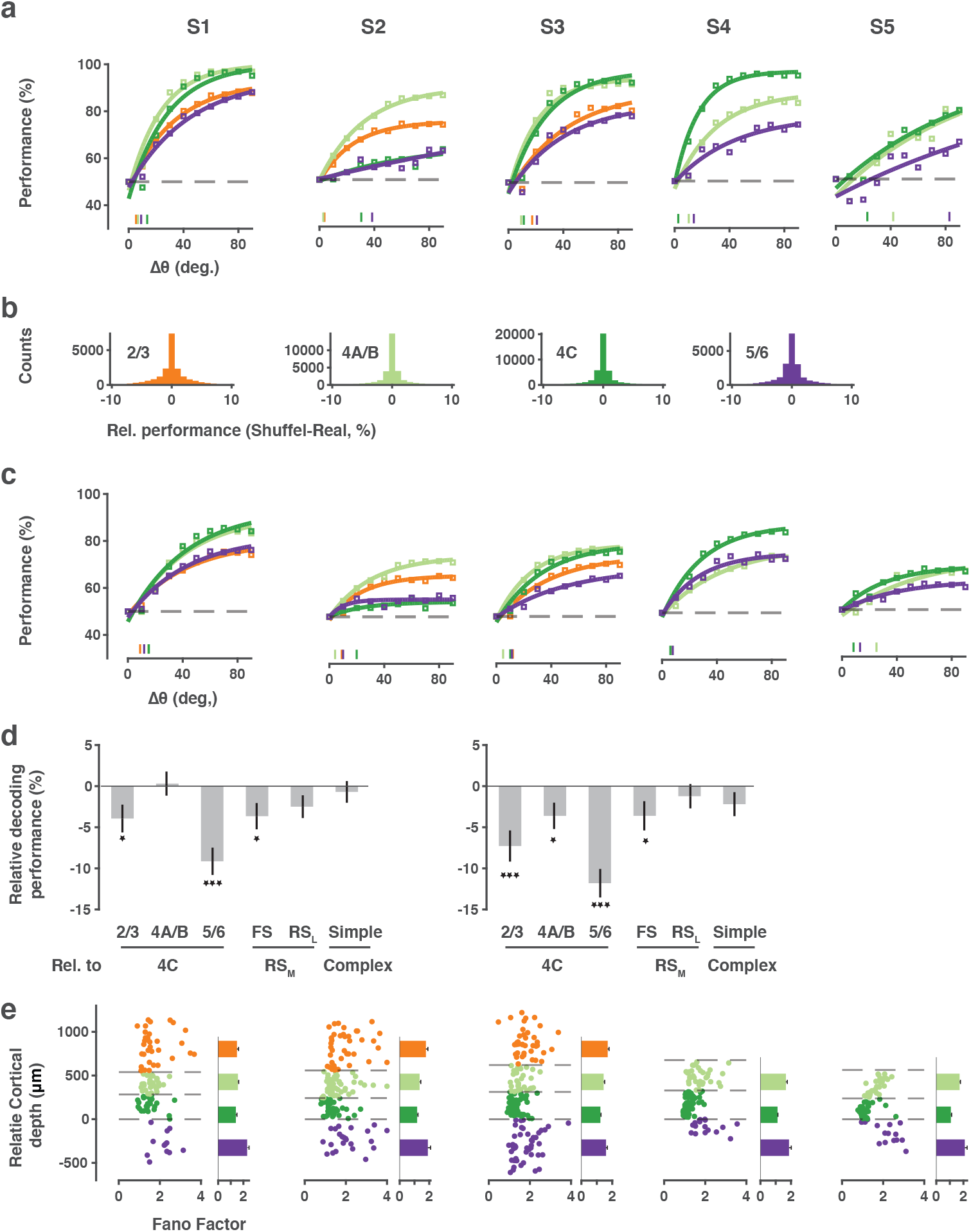
Trial-shuffled and single-neuron decoding of orientation. **a,** Performance of orientation decoders within different laminar compartments using shuffled trials. As in Fig. 6A, each point shows the average decoder performance using the activity of 10 neurons within each subpopulation for 9 pairwise orientation discriminations, but with shuffled trials. **b,** Distribution of differences in decoding performance between shuffled and unshuffled trials for each laminar compartment. Data from all pairwise decoding performances are shown. **c,** Mean single-neuron decoding performance within different laminar compartments. **d,** GLM coefficients for different predictors of decoding performance using all 5 (left) or the 4 columnar recordings sessions (right). Error bars denote +/− S.E.M.. **e,** Fano factors for single neurons within different laminar compartments.

Given that single-neuron properties seemed to be contributing more to differences in decoding performance, we used the same decoder to discriminate orientation from the activity of single neurons within different laminar compartments (Fig. 7C) (Methods). As expected, the overall average performance of single-neuron decoders at discriminating orientation pairs was reduced compared to that of 10-neuron populations. However, the pattern of results was remarkably similar between the single and population-level analyses. Specifically, in columnar sessions, the suprathreshold performance of layer 4C single-neuron decoders significantly exceeded that of superficial (L2/3) and deep (L5/6) layers (4C vs 2/3: mean Δ% = 8.5 ± 3.0; 4C vs 5/6: mean Δ%=9.9 ± 1.2; *P*=.0312, Two-tailed Wilcoxon signed-rank test). Decoding performance for layer 4A/B neurons also exceeded that of superficial (L2/3) and deep (L5/6) layers (4A/B vs 2/3: mean Δ% = 8.2 ± 1.9; 4A/B vs 5/6: mean Δ%=6.9 ± 2.6; *P*=.0312, Two-tailed Wilcoxon signed-rank test), similar to the population-level comparisons. Thus, the properties of single neurons were sufficient to yield superior performance of layer 4 decoders. We observed a similar pattern of results when comparing performance using a multi-class decoder, and across spatial frequency (Fig. S5).

As previously described, layer 4 neurons differ from those in superficial and deep layers in other important ways. For example, we observed clear differences in the proportions of different waveform classes across the four laminar compartments (Fig. 2). Furthermore, as expected, layer 4C contained a larger proportion of simple cells (Fig. 3). Thus, it is possible that differences in the performance of single cell decoders was associated more with those properties than with layer. To examine this possibility, we built a Generalized Linear Model (GLM) to measure the influence of three factors: waveform class (RS_L_, RS_M_ and FS), functional class (simple or complex), and laminar compartment. The GLM was built from data obtained either from all five sessions or from the four columnar sessions (Fig. 7D). For both models, differences in single-neuron decoder performance were more robustly associated with laminar compartment than with waveform or functional class. In neither case were performance differences associated with the functional class of neurons; simple and complex cell decoders performed equally well (All sessions: Δcoeff. = −0.73%, p =.5532; columnar sessions: Δcoeff. = −1.89%, p =.1757). However, in both models, decoders made from the activity of RS_M_ neurons performed significantly greater than FS neurons (All sessions: Δcoeff. = −3.53%, P =.0185; columnar sessions: Δcoeff. = −3.53%, P =.0381), but were statistically equal to the performance of RS_L_ neurons (All sessions: Δcoeff. = −2.07%, P =.1112; columnar sessions: Δcoeff. = −0.91%, P =.5201). Performance differences associated with laminar compartments were generally larger. Layer 4C decoder performance exceeded that of layer 2/3 and 5/6 decoders in both models and by as much as 10% (All session: 2/3 - 4C = −4.34%, P =.0063; 5/6 - 4C = −8.66%, P = 3.76E^−08^; columnar sessions: 2/3 - 4C = −7.0%, P = 1.27E^−04^; 5/6 - 4C = −11.1%, P = 7.31E^−11^). Furthermore, in columnar sessions, layer 4C decoders significantly exceed the performance of layer 4A/B (4A/B - 4C = −4.29%, P =.0052). Thus, cortical layer was clearly a factor in determining the performance of single-neuron decoders.

Given the lack of substantial differences in the basic tuning measures between neurons across layers, e.g. orientation index and tuning bandwidth, it is surprising that single neuron decoding of layer 4 neurons exceeded that of superficial and deep neurons. However, these measures fail to fully capture the information available in sensory responses. In particular, these measures do not account for differences in the reliability of stimulus-driven responses between different neurons, for example differences in the Fano factor (FF) (Churchland et al., 2010; Churchland, Yu, Ryu, Santhanam, & Shenoy, 2006; Steinmetz & Moore, 2010). Across the full population of visually responsive neurons (N = 676), the relative performance of single-neuron decoders was negatively correlated with the FF of the corresponding neuron responses (r = −0.1033; p =.0072). Thus, we considered the possibility that layer 4C neuronal responses might be more reliable than those of superficial and deep neurons. Previous studies comparing the FFs of V1 neurons across layers yielded equivocal results (Gur & Snodderly, 2006; Hansen et al., 2012), perhaps due to comparatively small datasets. We compared the FFs of neurons within the different laminar compartments (Methods)(Fig. 7E). We found highly robust differences in the FF of visual responses across laminar compartments (2/3 (n=116): 1.87+0.07, 4A/B (n=179): 1.62+0.04, 4C (n=247): 1.28+0.02, 5/6 (n=119): 1.94+0.06; Kruskal-Wallis test, ***χ***^2^(3) =167.32, *P*=4.82E^−36^). Moreover, the FFs of layer 4C neurons were significantly lower than that of neurons in layers 2/3 (*P*=5.22E^−22^, Two-tailed Mann-Whitney U test), 5/6 (*P*=3.53E^−23^), and also layer 4A/ B (*P*=4.16E^−19^). The Fano factors of layer 4A/B neurons were also significantly lower than that of neurons in layers 2/3 (*P*=4.30E^−3^), 5/6 (*P*=6.30E^−5^). These differences were not a result of differences in firing rate across layers (Fig. S6A) and they were present across all stimulus orientations (Fig. S6B). Moreover, these differences were largely independently of cell type (Fig. S6C). Thus, the greater performance of layer 4C neurons was associated with larger reliability in single neuron responses. Given that orientation selectivity indices and tuning bandwidths of layer 4C neurons were largely comparable to those in other layers (Fig. 5E-F), it is likely that the superior orientation decoding in 4C is due at least in part to greater response reliability of 4C neurons.

## Discussion

We studied the visual activity of large populations of neurons distributed across layers of primate V1 using high-density Neuropixels probes. The high capacity of the probes yielded single-neuron recordings from substantial numbers of nearby cells located within the same laminar compartments of single cortical columns, thereby facilitating robust comparisons between different subpopulations of neurons in single experiments. These comparisons revealed myriad differences in the functional properties of neurons across layers as well as between neurons of different putative cell types. In the latter case, putative inhibitory (FS) neurons, subclasses of putative excitatory (RS) neurons were distributed differently across the cortical depths and within laminar compartments, consistent with previous anatomical evidence (Fitzpatrick, Lund, Schmechel, & Towles, 1987; Hendry, Schwark, Jones, & Yan, 1987). In the former case, several noteworthy functional differences between neuronal subpopulations were observed. First, we observed robust differences in synchronized and correlated activity between neuron pairs within and across laminar compartments. Both synchrony and noise correlations were dramatically reduced among pairs involving layer 4C neurons, compared to all other laminar pairs. This observation expands on an earlier report of surprisingly low noise correlations in layer 4C (Hansen et al., 2012). Our results revealed a clear decrement in synchrony coinciding with the superficial and deep borders of 4Cα and 4Cβ, respectively (Fig. 3E). Although lower synchrony and noise correlations can be a result of correspondingly low shared inputs, this is an unlikely basis for the input layers of V1 given that neurons there share inputs and are highly connected. Instead, it may be that the reduced level of coordinated activity in layer 4C results from a greater balance of excitatory and inhibitory inputs, which has been shown to limit correlations in highly connected neurons (Renart et al., 2010). Indeed, this is consistent with the relatively high ratio of FS neurons to RS neurons we observed within layer 4C (Fig. 2C).

In contrast to synchrony and noise correlations, signal correlations among layer 4C neurons were elevated compared to that of other laminar compartments. Furthermore, the relationship between synchrony and signal correlations within layer 4C differed dramatically from that of all other compartments; synchrony remained very low even for neurons with high signal correlations, and thus similar tuning properties. In addition, we found that both noise and signal correlations were generally lower among FS neurons than among RS cells. Notably, this latter observation directly contrasts with the pattern reported in mouse visual cortex in which greater correlations, both signal and noise, have been observed among inhibitory neurons, including parvalbumin+ (Hofer et al., 2011) and other inhibitory subtypes (de Vries et al., 2020). This contrast suggests a distinct difference between the functional microcircuitry of mouse and primate V1.

In addition, single recording sessions could reveal a number of well-known differences in the response properties of neurons across cortical layers, including higher firing rates, but lower proportions of complex cells or direction-selective neurons in layer 4C. However, as with many previous studies, comparisons of orientation selectivity across laminae using standard measures yielded equivocal results in terms of identifying clear differences in orientation selectivity between layers. We therefore used a decoder to measure the orientation discrimination achievable from the activity of populations within different laminar compartments. Surprisingly, we found that in columnar recording sessions, decoder performance in layer 4C was superior to that of superficial and deep layers. This is the first observation of an unambiguous difference in orientation discrimination between neurons spanning different layers of the same V1 column. Importantly, the superior orientation discrimination from layer 4C activity was not dependent upon differences in correlated variability observed between laminar compartments, as might be predicted (Hansen et al., 2012). Instead, single-neuron decoding yielded an identical pattern of results, with orientation decoding from layer 4C neurons exceeding the performance of superficial and deep layer neurons. In addition, the superior orientation discrimination was associated with reduced response variability in layer 4C neuronal responses. This result not only contrasts with the classic view that orientation selectivity is largely absent among neurons in layer 4C of primate V1, particularly 4Cβ (Blasdel & Fitzpatrick, 1984; Livingstone & Hubel, 1984), and confirms earlier evidence of clear orientation tuning in 4C (Hawken & Parker, 1984; Livingstone & Hubel, 1984; Ringach et al., 2002; Schiller et al., 1976), but it demonstrates that when comparing neurons within the same column, the fidelity of orientation information is at its peak in the output of layer 4C neurons.

The visual system of the macaque monkey has proven to be remarkably similar to that of the human and is thus an ideal model. However, in contrast to simpler model systems, extracting circuit-level information from studies of the nonhuman primate visual system has proven particularly challenging given the limited arsenal of appropriate tools. One key shortcoming of past neurophysiological studies in nonhuman primates is their relative inability to capture the diversity of neural signals present within both local and distributed populations of neurons in simultaneous recordings. Neurophysiological studies within the primate visual system have largely involved successive recordings from individual neurons, or small numbers of neurons using conventional single-electrodes, or low-channel count linear arrays. From such data, neuronal properties are studied in aggregated datasets of recordings accumulated across multiple sessions. As a result, direct comparisons between subpopulations of neurons within local circuits, e.g. within single cortical columns, are less than ideal. Although many studies employing implanted arrays have yielded datasets from ~100s simultaneously recorded neurons, particularly within the motor system (Churchland et al., 2012; Wannig, Stanisor, & Roelfsema, 2011), such recordings can only be achieved within surface (and flat) cortical areas, and importantly, tend to restrict sampling of neurons at a fixed depth. A number of recent studies have demonstrated the advantages of recently developed high-density silicon probes, particularly Neuropixels probes, in capturing the properties and dynamics of large populations of local and distributed neurons (Jun et al., 2017; Steinmetz, Koch, Harris, & Carandini, 2018; Steinmetz et al., 2019). In these first high-density recordings of primate V1, we have shown the value of such an approach in revealing the major properties of neurons comprising neocortical columns, a fundamental unit of neocortical circuitry.

## Methods

### Electrophysiological Recordings

Anesthetized recordings were conducted in 2 adult male macaques (M1, x kg, M2 x kg. All experimental procedures were in accordance with National Institutes of Health Guide for the Care and Use of Laboratory Animals, the Society for Neuroscience Guidelines and Policies, and Stanford University Animal Care and Use Committee. Prior to each recording session, treatment with dexamethasone phosphate (2 mg per 24 h) was instituted 24 h to reduce cerebral edema. After administration of ketamine HCl (10 mg per kilogram body weight, intramuscularly), monkeys were ventilated with 0.5% isoflurane in a 1:1 mixture of N2O and O2 to maintain general anesthesia. Electrocardiogram, respiratory rate, body temperature, blood oxygenation, end-tidal CO2, urine output and inspired/expired concentrations of anesthetic gases were monitored continuously. Normal saline was given intravenously at a variable rate to maintain adequate urine output. After a cycloplegic agent was administered, the eyes were focused with contact lenses on a CRT monitor. Vecuronium bromide (60 μg/kg/h) was infused to prevent eye movements.

With the anesthetized monkey in the stereotaxic frame, an occipital craniotomy was performed over the opercular surface of V1. The dura was reflected to expose a small (~3 mm^2^) patch of cortex. Next, a region relatively devoid of large surface vessels was selected for implantation, and the Neuropixels probe was inserted with the aid of a surgical microscope. Given the width of the probe (70 um x 20 um), insertion of it into the cortex sometimes required multiple attempts if it flexed upon contacting the pia. The junction of the probe tip and the pia could be visualized via the (Zeiss) surgical scope and the relaxation of pia dimpling was used to indicate penetration, after which the probe was lowered at least 3-4 mm. Prior to probe insertion, it was dipped in a solution of the DiI derivative FM1-43FX (Molecular Probes, Inc) for subsequent histological visualization of the electrode track.

Given the length of the probe (1 cm), and the complete distribution of electrode contacts throughout its length, recordings could be made either in the opercular surface cortex (M1) or within the underlying calcarine sulcus (M2), by selecting a subset of contiguous set of active contacts (n = 384) from the total number (n=986). Receptive fields (RFs) from online multi-unit activity were localized on the display using at least one eye. RF eccentricities were ~ 4-6° (M1) and ~ 6-10° (M2). Recordings were made at 1 to 3 sites in one hemisphere of each monkey. At the end of the experiment, monkeys were euthanized with pentobarbital (150 mg kg−1) and perfused with normal saline followed by 1 liter of 1% (wt/vol) paraformaldehyde in 0.1 M phosphate buffer, pH 7.4.

### Visual Stimulation

Visual stimuli were presented on a LCD monitor NEC-4010 (Dimensions= 88.5 (H)* 49.7 (V) cm, pixels=1360 * 768, frame rate= 60 Hz) positioned 114 cm from the monkey. Stimuli consisted of circular drifting Gabor gratings (2 deg./sec., 100% Michelson contrast) positioned within the joint RFs of recorded neurons monocularly. Gratings drifted in 36 different directions between 0 to 360° in 10° steps in a pseudorandom order. Four spatial frequencies (0.5, 1, 2, 4 cycle/deg.) were tested and optimal SFs were determined offline for data analysis. The stimulus in each condition was presented for 1s and repeated 5 or 10 times. A blank screen with equal luminance to the Gabor patch was presented for 0.25s during the stimulus interval.

### Layer Assignment

The laminar location of our recording sites was estimated based on a combination of functional analysis and histology results. For each recording, we first performed the current source density (CSD) analysis on the stimulus-triggered average of LFP. LFP signals recorded from each 4 neighboring channels were averaged and realigned to the onset of visual stimulus. CSD was estimated as the second-order derivatives of signals along the probe axis using the common five-point formula (Nicholson & Freeman, 1975). The result was then smoothed across space (σ = 120 μm) to reduce the artifact caused by varied electrode impedance. We located the lower boundary of the major sink (the reversal point of sink and source) as the border between layer 4C and layer 5/6. Based on this anchor point, we assign other laminar compartment borders using the histological estimates.

### Waveform Classification

We classified different waveform types based on the trough-peak duration of the spike template of all recorded neurons (including the non-visual responsive cells). For each spike template, we measured the trough-peak duration by subtracting the time of the maximum value from the time of the minimum value. In some cases, the peak before the trough has a greater amplitude than the peak following the trough, leading to a negative duration value. We defined this group of neurons as putative “axonal” units (Schomburg et al., 2012). For the rest of waveforms, we divided them into 2 classes based on the peak-trough duration, referred as “fast-spiking” (≤ 200 μs) and “regular-spiking” (> 200 μs) units. The classification boundary of duration was chosen in consistent with previous studies (Mitchell et al., 2007). To investigate the heterogeneity of waveforms in the regular-spiking class, we further divided these waveforms into “regular-spiking (medium)” and “regular-spiking (long)” units using an arbitrarily selected boundary (300 μs).

### Cross-correlograms

The pairwise cross-correlogram (CCG) was calculated based on spike trains of simultaneously recorded neurons (Perkel, Gerstein, & Moore, 1967a). To make sure that our analysis was not affected by the instationary transient response, we considered only the spiking activity during the 0.4 ~ 0.6 s period of each visual stimulation, and computed the CCG for each pair (j,k) as a function of time lag τ:

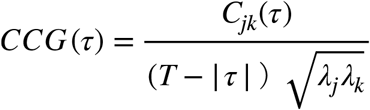

where *C_jk_*(*τ*) is the auto-/cross- correlation calculated on spike train data from trials of the same condition; T is the total time length of the spike train (0.2 s); *λ_j_* and *λ_k_* are the mean firing rate of neuron j and neuron k, respectively. The divisor terms serve to normalize the result so that CCG remains relatively constant with varied τ and firing rates. To further eliminate the component in CCG that is attributed to the stimulus-locked activity, we computed a shift- (or shuffle-) predictor based on non-simultaneous responses (i.e., post-stimulus time histograms, PSTH):

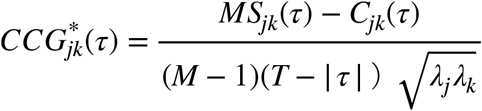

Where M is the trial number; *S_jk_*(*τ*) denotes the auto-/cross- correlation calculated on PSTHs of neuron j and neuron k. For every pair of simultaneously recorded neurons, a CCG was computed for each unique condition, shift-corrected (CCG minus CCG^*^), and then averaged among all conditions that yielded non-zero visual responses. Synchrony between a pair of neurons was defined as the peak (maximum) value of the shift-corrected CCG function. Direction of signal transfer was estimated using the temporal delay (τ) that corresponds to the CCG peak.

To be comparable with the CCG measurements, noise correlation and signal correlation were also calculated using spiking activity during the 0.4 ~ 0.6 s period of each visual stimulation for each pair of neurons. As commonly defined, noise correlation between two neurons was estimated with the correlation coefficient of their spike counts across repetitive trials under the same experimental conditions and averaged over conditions. Signal correlation was estimated with the correlation coefficient of their mean spike counts across different experimental conditions.

### Single neuron properties

To characterize neuronal properties, the evoked activity was assessed using mean firing rate (spikes/sec) over the whole stimulus presentation period, offset by response latency delay. Only responses to the preferred spatial frequency were selected. The maximum firing rate was the neuron’s response to the preferred drifting orientation and direction. Modulation ratio was defined as F1/F0, where F1 and F0 are the amplitude of the first harmonic at the temporal frequency of drifting grating and constant component of the Fourier spectrum to the neuron’s response to preferred orientation. Direction selectivity (Direction Index, DI) was determined as the response to preferred orientation and drift direction minus the response to preferred orientation but opposite drift direction, divided by the sum of these two responses (Swindale, 1998). Orientation selectivity (Orientation Index, OI) was determined as the response to preferred orientation minus the response to orthogonal orientation, divided by the sum of these two responses (Swindale, 1998). To estimate the orientation tuning bandwidth, the orientation tuning responses were first smoothed with a hanning window (half width at half height of 20^°^), and then fitted with a von-Mises function (Swindale, 1998)

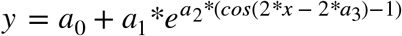

Only neurons that were well fit by the function (R^2^>0.7) were included in the bandwidth analysis. The locations of the peak of the fitted curves were determined. The two orientations closest to the peak on either side of the tuning curve where responses dropped to 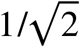 of the peak response were then estimated (Schiller et al., 1976). Bandwidth was defined as the half of the differences between the two orientations. If the response around the peak never went below the response criteria, the tuning bandwidth was defined as 180^°^.

Fano factor were computed to assess individual neuron’s response variability. For each stimulus condition and 100ms time window bin, spikes occurrences were counted for all the trial repetitions. And Fano factor was defined as the spike counts variance across trials divided by spike counts mean. And then Fano factors generated from the whole stimulus presentation period to all the stimulus conditions were then averaged to determine the Fano factor for that neuron.

### LDA Decoding

Linear discriminant analysis (LDA) decoders were performed to quantify the ability of neuronal populations in encoding stimulus orientations. The analysis was performed using Matlab function “Classify” and the “diaglinear” option (Fisher, 1936; Mendoza-Halliday & Martinez-Trujillo, 2017). For each recording session, multiple LDA decoders were built from each neuronal subpopulation’s responses to each pair of stimulus orientations. Neuronal subpopulations consisted of a fixed number of 10 nearby single neurons, and the depth of neuron in the center was used to assign this subpopulation to corresponding laminar compartment. Responses were calculated as the mean spike counts within the whole visual stimulation period (50 ms to 1050 ms after stimulus onset, considering response latency). Each orientation pair consisted of the preferred orientation of the 10-neuron subpopulation, and another orientation at varying differences from the preferred (Δθ=10°, 20°, …90°). Only trials tested with gratings of optimal spatial frequency of the subpopulation were selected, which resulted in 20-40 trials tested for a given orientation pair. A leave-2-out cross validation was performed using a repeated random subsample technique (All except a random selection of two trials for training, remaining two trials for testing, ~200 repetitions). Decoding performance was defined as the percentage of correct classification of all the repetitions. For each laminar compartment, decoding performance of all the subpopulations within it were averaged, and plotted as a function of the difference between orientations. It was then fitted with a saturation function: f~a*e^−b*x^ + c using non-linear least-square option in Matlab.

Instead of a fixed number of neurons, Neuronal Dropping Curves (NDCs) were generated using population decoders with varied neuronal subpopulation size. For a recording session, we systematically selected n (n=1, 2, …) single neurons located within each laminar compartment. Each selection for a given n consisted of a different combination of neurons and was repeated for a maximum of 200 times. The orientation pairs to discriminate consisted of the preferred orientation (θpref) for the whole recording session, and one of 8 other orientations (θ_pref_ ± 20°, θ_pref_ ± 40°, θ_pref_ ± 60°, θ_pref_ ± 80°). Decoding performance for these 8 orientation pairs were later averaged.

Decoding sensitivity was determined for each neuronal subpopulation at each orientation. The orientation pairs to discriminate were extended to all pairwise combinations of 18 tested orientations, which resulted in 153 pairs of orientations. From the decoder performance at discriminating a given orientation from orientations with varying differences (Δθ=10°, 20°, …90°), a minimum difference Δθ_min_ was interpolated so that decoder performance could reach an arbitrary threshold level (60% in this case). Sensitivity was simply the inverse of Δθ_min_. A decoder with a high sensitivity can robustly discriminate orientation pairs with small differences.

Shuffled population decoders were built almost the same way, except that trials of all the 10 neurons in a subpopulation were randomly shuffled independently. The relative contribution of shuffling on decoding was assessed by simply computing the difference in decoding performance between trial shuffled decoder and real data decoder. Decoding to all the pairwise orientation discrimination was included.

Single neuron decoders were also built almost the same way as the population decoder, except that instead of using 10 neurons, only one neuron was used. To quantify the influences of various factors on the decoding performance, Generalized Linear Model (GLM) was built. Decoding performance ~ Constant + Laminar Compartment + waveform class (RS_L_, RS_M_ and FS) + functional class (simple or complex).

## Acknowledgments

We thank Tim Harris and Karel Svoboda for providing the Neuropixel probes, Jonathan C. Horton for extensive help with the recordings and histology, Jonathan Nassi for helpful comments on the results and interpretations, and Shellie Hyde for technical assistance. This work was supported by NIH Grant EY014924.

---

**Supplementary table 1.** Density of FS waveforms:

|  | S01 | S02 | S03 | S04 | S05 | Mean |
| --- | --- | --- | --- | --- | --- | --- |
| 2/3 | 2.17 | 3.22 | 4.28 |  |  | 3.23 |
| 4A/B | 8.98 | 4.87 | 12.03 | 4.87 | 8.01 | 7.75 |
| 4C | 4.24 | 5.77 | 16.53 | 14.33 | 11.76 | 10.53 |
| 5/6 | 1.60 | 3.77 | 3.89 | 4.28 | 3.13 | 3.33 |

**Density of RS waveforms:**
|  | S01 | S02 | S03 | S04 | S05 | Mean |
| --- | --- | --- | --- | --- | --- | --- |
| 2/3 | 7.96 | 14.57 | 9.88 |  |  | 10.80 |
| 4A/B | 10.55 | 15.58 | 18.67 | 20.92 | 14.10 | 15.96 |
| 4C | 10.25 | 27.88 | 21.07 | 26.83 | 18.49 | 20.90 |
| 5/6 | 1.96 | 10.68 | 10.25 | 8.82 | 9.92 | 8.32 |

**Supplementary table 2.**
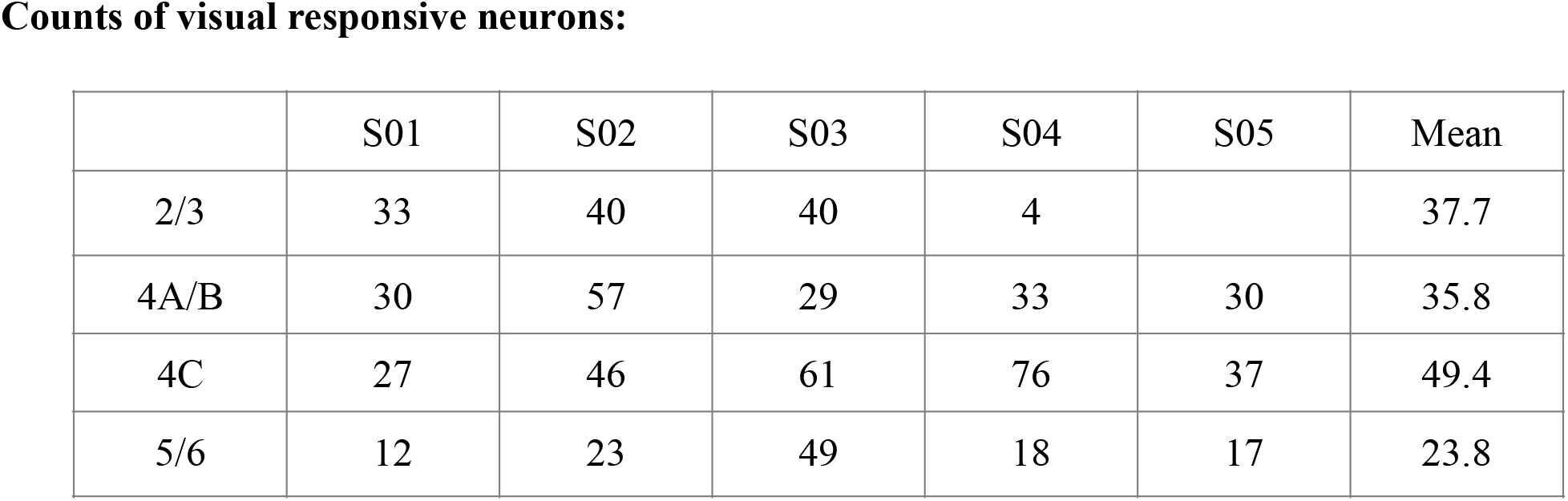

**Supplementary table 3.** Comparison of synchrony for different waveform types (table shows t-test p values)

|  | FS | RS<sub>m</sub> | RS<sub>l</sub> |
| --- | --- | --- | --- |
| FS | 1 | 0.000002 | 0.000015 |
| RS<sub>m</sub> | 0.000002 | 1 | 0.148420 |
| RS<sub>l</sub> | 0.000015 | 0.148420 | 1 |

**Comparison of synchrony for different laminar compartments.**
|  | 5/6 | 4C | 4A/B | 2/3 |
| --- | --- | --- | --- | --- |
| 5/6 | 1 | 1.822276e-56 | 2.564906e-02 | 1.446454e-06 |
| 4C | 1.822276e-56 | 1 | 2.369309e-104 | 3.235028e-91 |
| 4A/B | 2.564906e-02 | 2.369309e-104 | 1 | 1.779898e-05 |
| 2/3 | 1.446454e-06 | 3.235028e-91 | 1.779898e-05 | 1 |

**Supplementary table 4.** Comparison of noise correlation for different waveform types.

|  | FS | RS<sub>m</sub> | RS<sub>l</sub> |
| --- | --- | --- | --- |
| FS | 1 | 5.863282e-39 | 3.925978e-56 |
| RS<sub>m</sub> | 5.863282e-39 | 1 | 7.170126e-03 |
| RS<sub>l</sub> | 3.925978e-56 | 7.170126e-03 | 1 |

**Comparison of noise correlation for different laminar compartments.**
|  | 5/6 | 4C | 4A/B | 2/3 |
| --- | --- | --- | --- | --- |
| 5/6 | 1 | 6.003955e-08 | 1.483738e-01 | 4.260968e-03 |
| 4C | 6.003955e-08 | 1 | 1.879479e-20 | 8.791248e-26 |
| 4A/B | 1.483738e-01 | 1.879479e-20 | 1 | 6.064193e-02 |
| 2/3 | 4.260968e-03 | 8.791248e-26 | 6.064193e-02 | 1 |

**Supplementary table 5.**
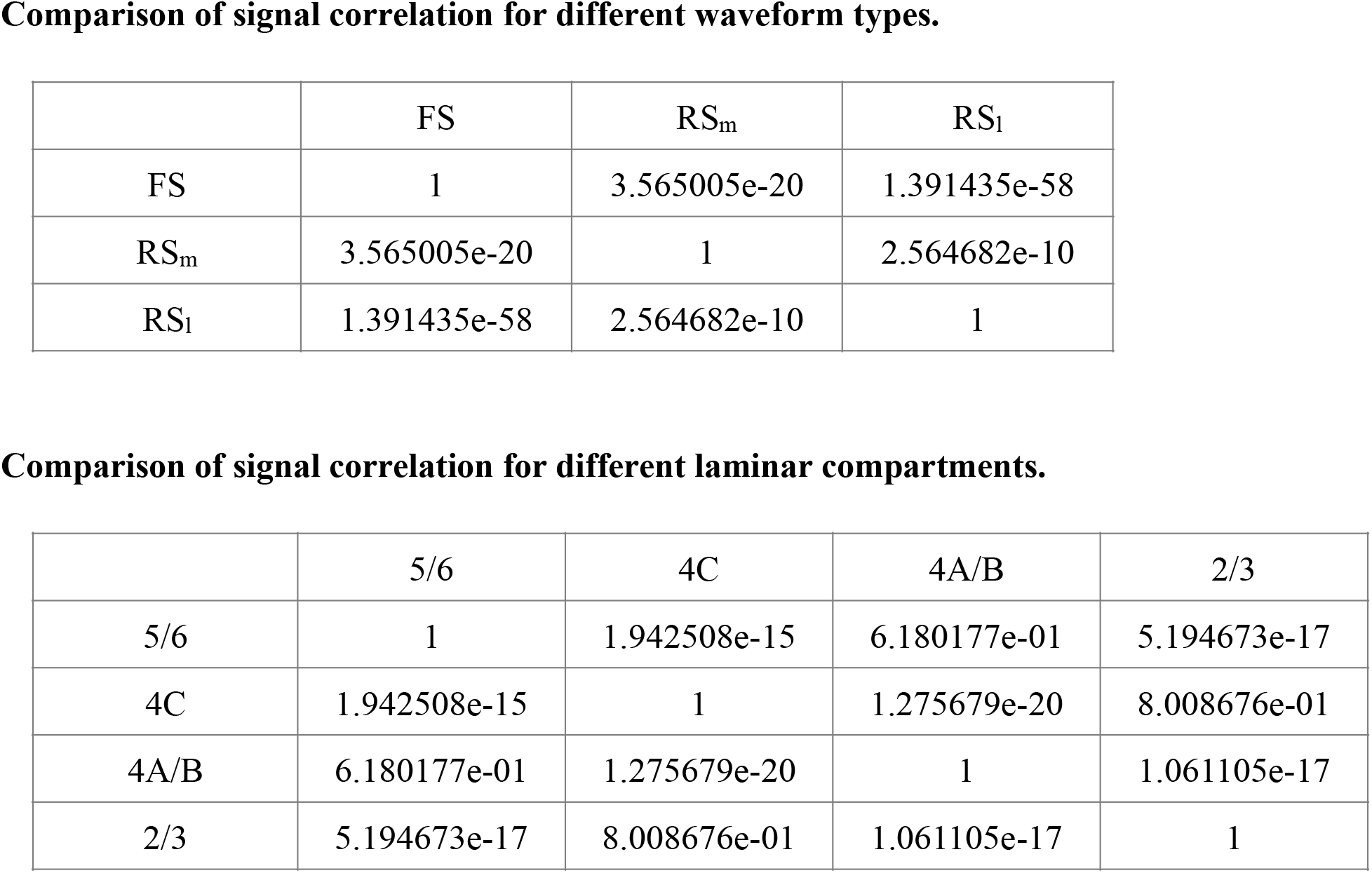

**Fig S1.**
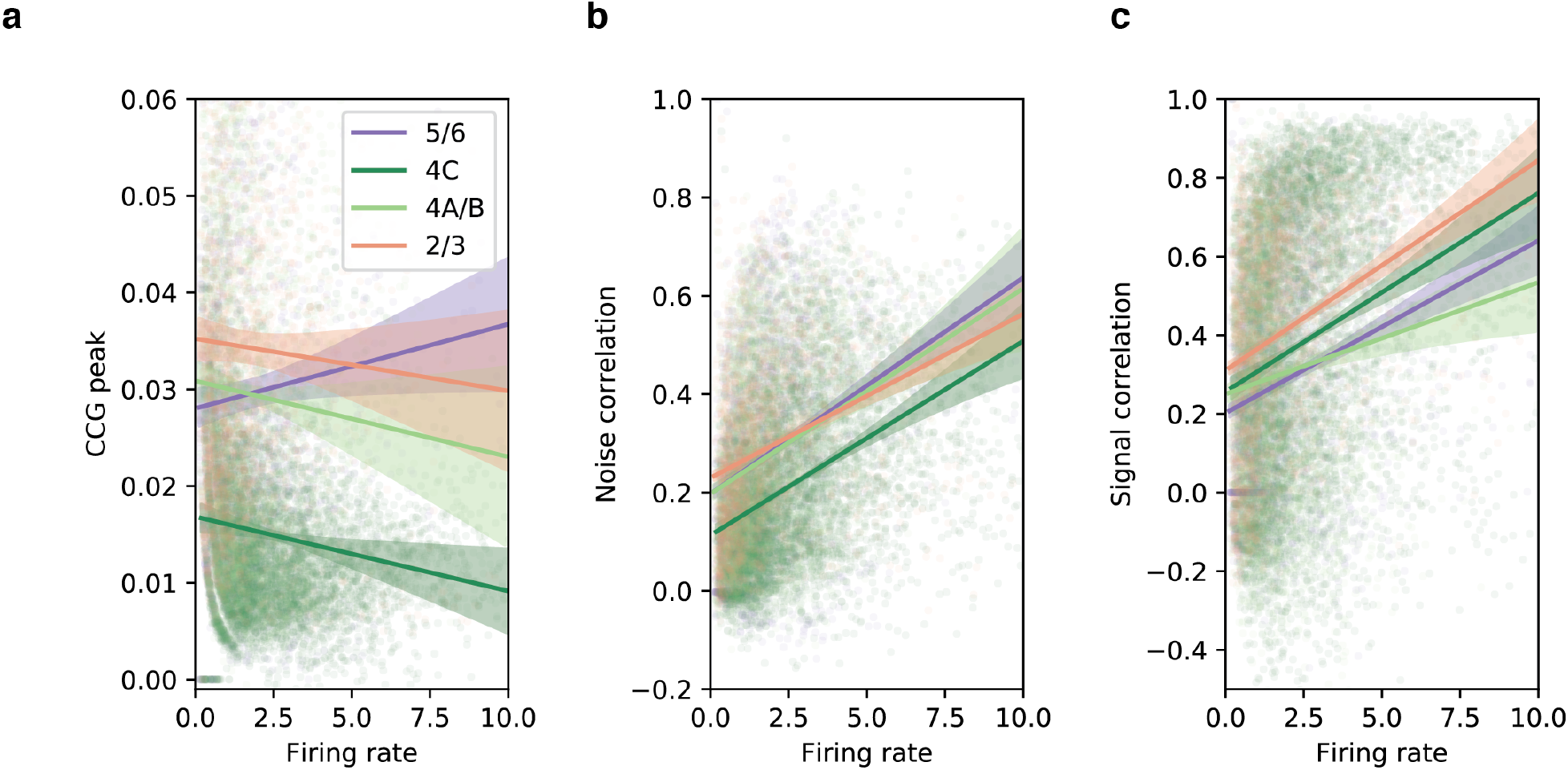
Correlated activity across layers as a function of firing rate. Scatter plots show the CCG peak, noise correlation, and signal correlation against the geometric mean of firing rate for pairs of neurons, respectively. Colors represent layer compartments where neuron pairs belong. Lines denote the linear fit for each layer with shaded areas representing the 95% confidence interval of fitting.

**Fig S2.**
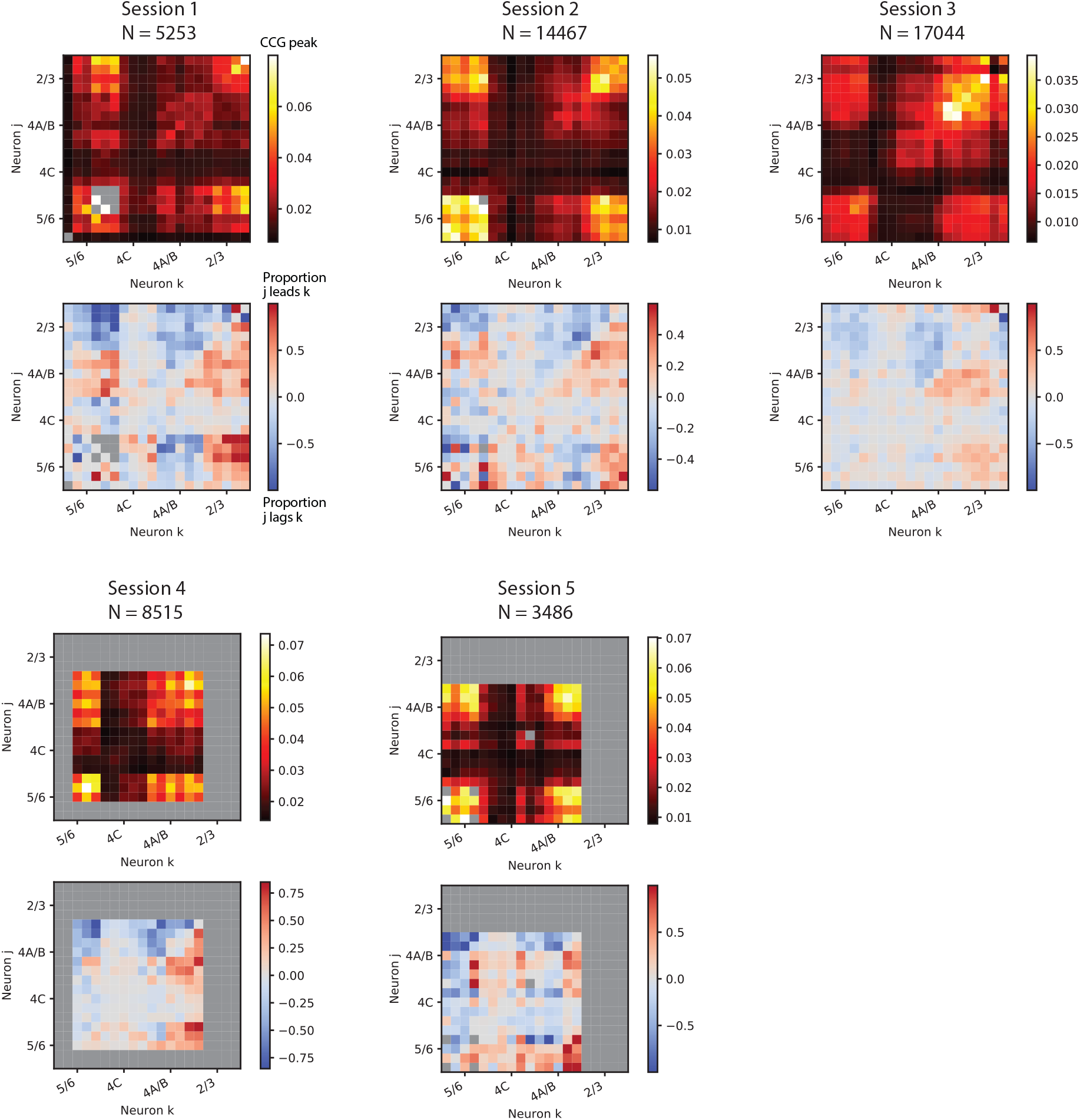
Cross-correlations in neuronal activity across V1 layers for all individual sessions. The CCG peak and the CCG direction matrices are shown for each of the 5 sessions. Plots follow the same color scheme used in Figure 3.

**Fig S3.**
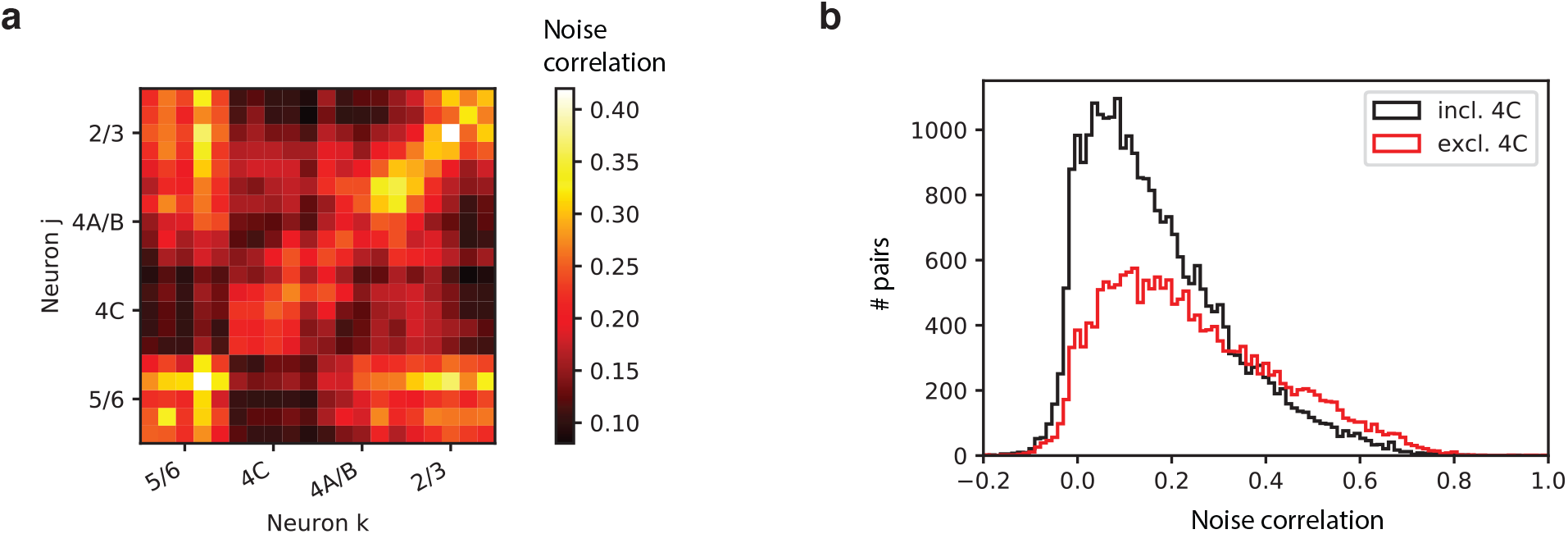
Noise correlation across V1 layers. **a,** Matrix of noise correlation across cortical depth for all sessions combined. **b,** A comparison between distributions of noise correlation for pairs that include vs. exclude layer 4C neurons.

**Fig S4.**
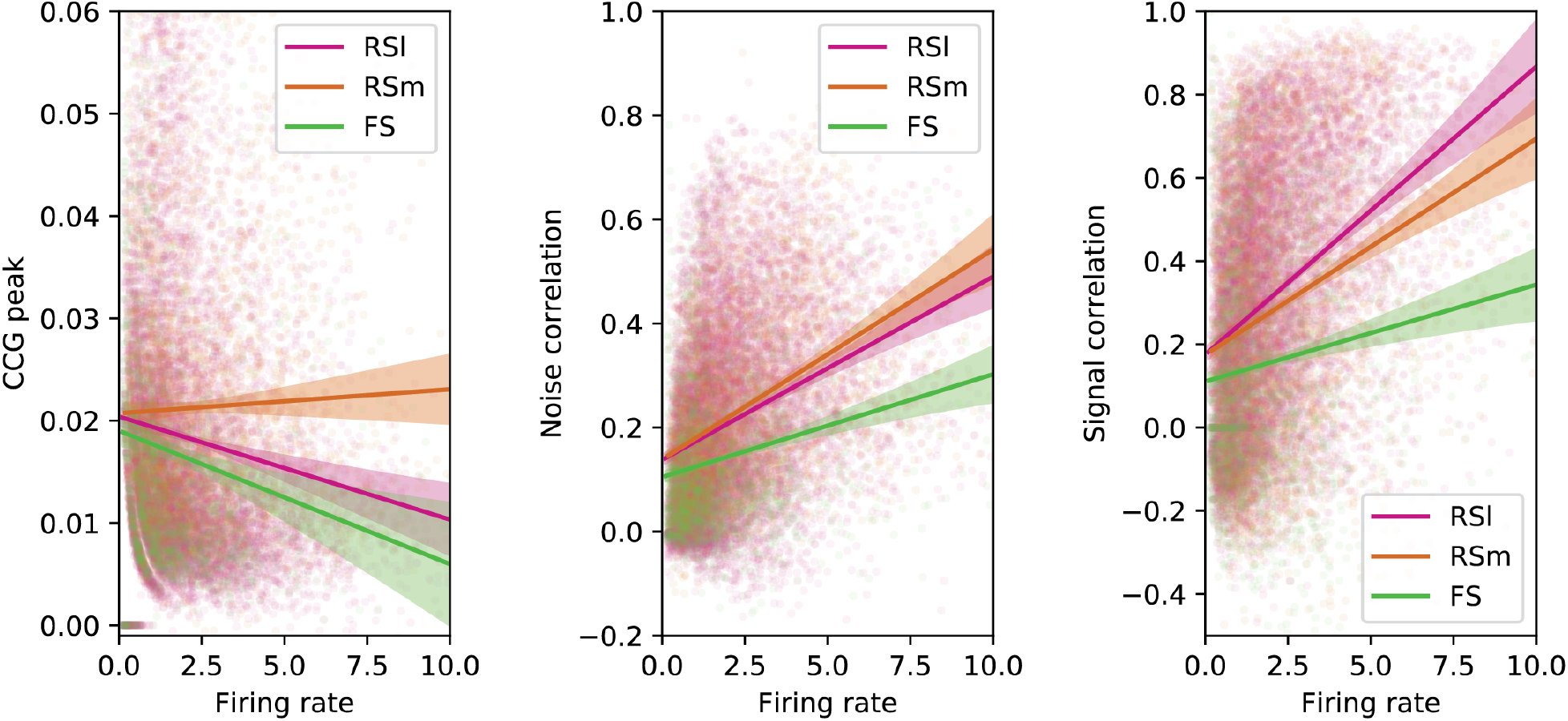
Correlated activity across waveform types as a function of firing rate. Scatter plots show the CCG peak, noise correlation, and signal correlation against the geometric mean of firing rate for pairs of neurons, respectively. Colors represent the neuron pairs’ waveform types. Lines denote the linear fit for each waveform type combination with shaded areas representing the 95% confidence interval of fitting.

**Fig S5.**
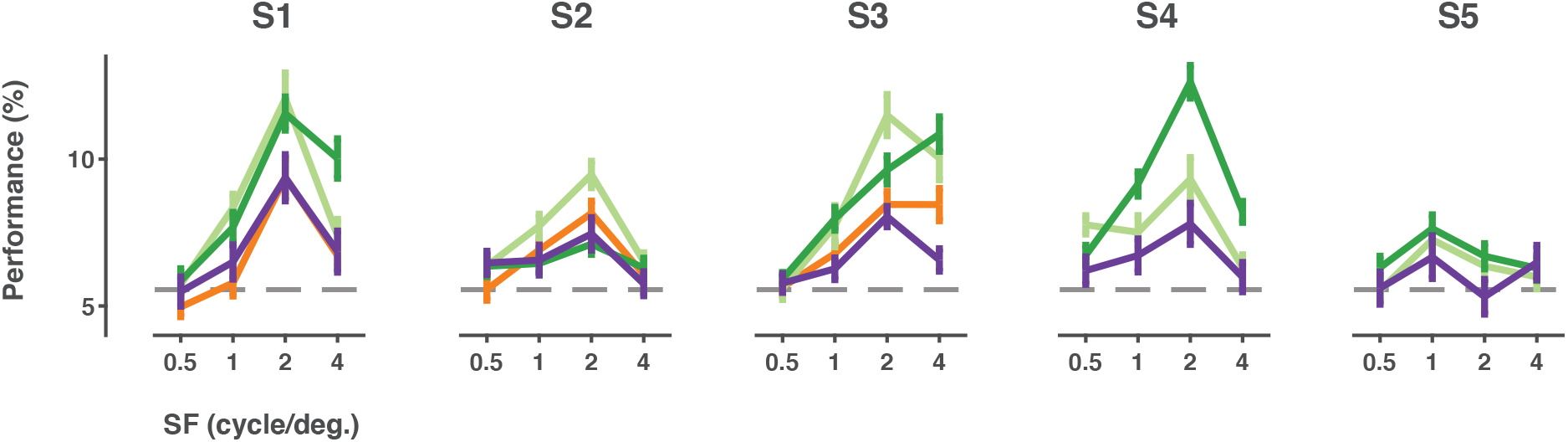
Single neuron, multi-class decoding of orientation across stimuli spatial frequencies. Decoders were trained using activity of single neuron to discriminate all 18 different stimuli orientations tested for each spatial frequency. Each point shows the mean decoder performance from neurons within each laminar compartment. Different colored lines donate different laminar compartment. Error bars denote +/− S.E.M. Results from each recording session are shown separately.

**Fig S6.**
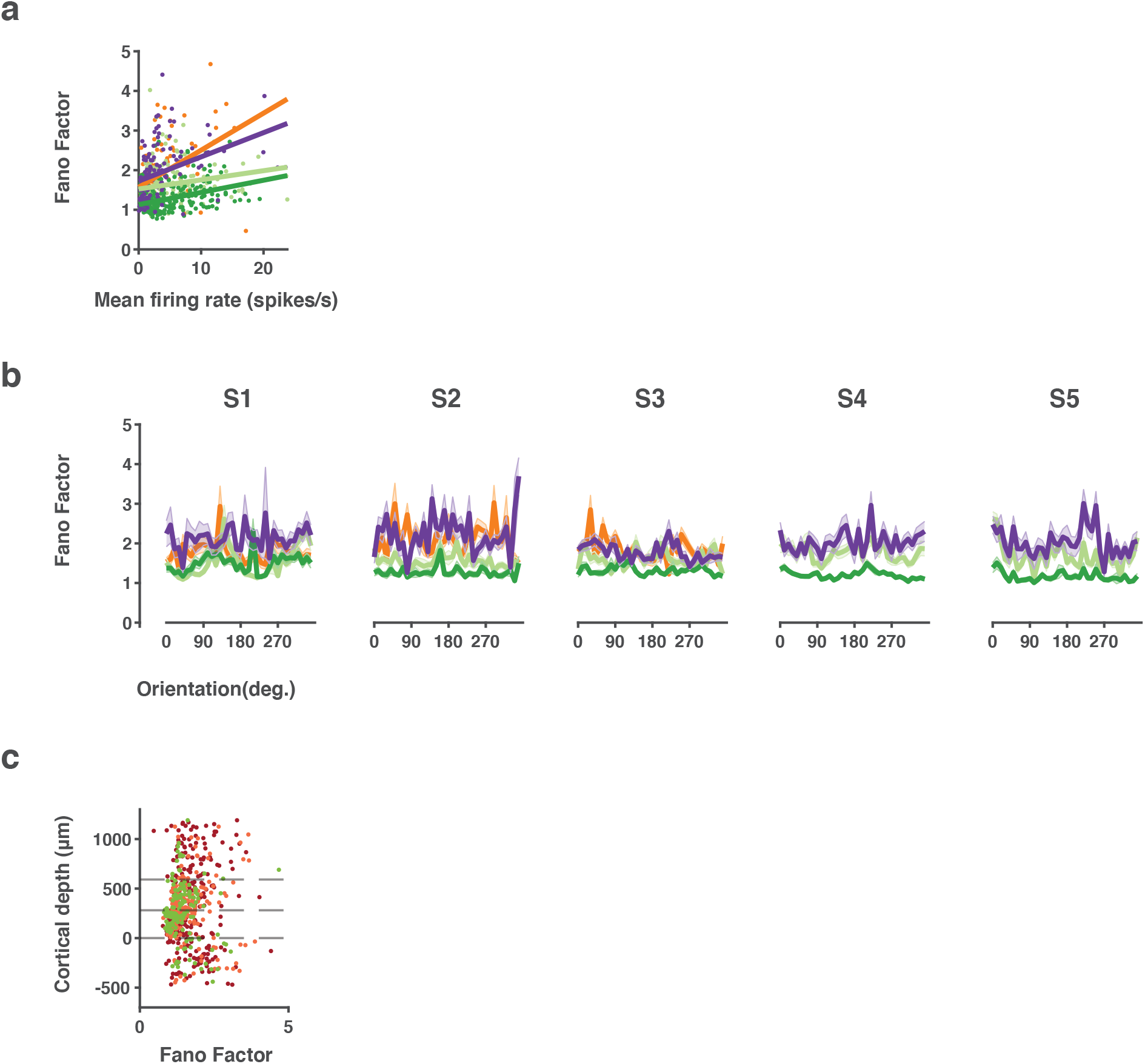
V1 neuronal response variability across laminar. **a,** Scatter plots showing Fano factor against mean firing rate for all neurons across different laminar compartment. Each point represents a single neuron, with different colors representing the corresponding laminar compartment it’s located in. Lines denote linear fits for each compartment. Results from all 5 recording sessions are combined. **b,** Fano factor across different stimuli conditions. Each point shows the mean Fano factor of neurons in each laminar compartment to each of the 18 different stimuli orientations. Error bars denote +/− S.E.M. Results from each recording session are shown separately. **c,** Laminar distribution of Fano factor across different cell types. Each point shows the Fano factor of each neuron plotted against its relative cortical depth. Different colors represent different waveform cell types. Results from all recording sessions are combined.

